# AI-guided discovery of natural pocket proteins for portable small-molecule sensing

**DOI:** 10.64898/2026.10.22.760851

**Authors:** Liping Huang, Yihui Yang, Changzhi Jiang, Hongli Fan, Yuhong Li, Yiyi Zhang, Yuxue Zhang, Deyi Li, Hanlin Zhou, Youqian Chen, Wenjun Hu, Jianguo Guan, Xiangrong Liu, Chunmeng Sun, Gang L. Liu

## Abstract

Recognition-element development remains an important practical challenge in small-molecule biosensing. Here, we present Dry-Wet Closed-Loop AI-driven Sensing Strategy (DWCLASS), an AI-guided workflow that combines screening of natural pocket proteins, molecular dynamics simulations, experimental binding assessment, and portable metasurface plasmonic sensing. Using thiamine (vitamin B1, thiamine) and 25-hydroxyvitamin D3 [25(OH)D_3_] as model analytes, the workflow identified binding responses in 7 of 16 tested analyte–protein pairs (43.8%). Selected proteins were incorporated into a competitive MetaSPR assay with gold nanoparticle amplification and acoustic mixing. The portable device provided a 240 s measurement step for prepared samples. Serum LODs were 2.98 ppt for thiamine and 3.82 ppt for 25(OH)D₃. Preliminary measurements of human serum specimens were compared with commercial ELISA to explore performance in biological samples. This study establishes an integrated route from AI-assisted screening of natural proteins to portable assay development for two chemically distinct vitamin analytes. Applicability to additional targets and clinical reference accuracy remain to be established.

## 1. Introduction

Small molecules are the most abundant class of biomarkers in human physiology, regulating nearly all biological processes from metabolism to cell signaling^[1,2]^. Their dysregulation is associated with human diseases, including nutritional deficiencies, metabolic disorders, cancer, and cardiovascular disease^[3]^. Their measurement at the point of care could support timely biochemical assessment where access to centralized testing is limited. For example, thiamine deficiency is associated with neurological and cardiovascular complications^[4,5]^, while vitamin D status is important to bone and mineral metabolism^[6]^. Translating small-molecule measurements into portable formats requires both selective molecular recognition and a readout capable of resolving low-mass binding events. The development of recognition reagents that can operate effectively within a compact sensing interface therefore remains an important part of small-molecule biosensor design.

Existing analytical approaches offer complementary capabilities. LC–MS/MS provides molecular selectivity in laboratory analysis^[7–9]^, while ELISA and lateral-flow assays support established forms of affinity-based measurement^[10–12]^. Antibodies are effective recognition reagents for many small molecules, including targets for which picomolar affinities have been reported^[13]^. Their development for a new target can nevertheless require hapten design, immunization, and reagent optimization^[14,15]^. Natural binding proteins, aptamers, and computationally designed proteins offer additional routes to molecular recognition^[16–18]^. A central opportunity is to search existing protein structures systematically for binding pockets that can be repurposed as recognition elements for a specified small molecule.

In recent years, artificial intelligence (AI) has emerged as a promising tool to address the recognition element crisis^[19–21]^. Deep learning models have shown remarkable progress in predicting protein-ligand binding affinities, but their application has been almost exclusively limited to protein to small-molecule screening, identifying small molecules that bind to a given protein target^[22–24]^. The reverse task, identifying proteins that bind to a given small molecule, remains largely unexplored. Existing models suffer from poor generalization beyond their training distributions, and standalone virtual screening tools ignore protein conformational flexibility and real-world experimental conditions, resulting in hit rates as low as 17.5% even for state-of-the-art models like DrugCLIP^[25]^. Standalone AI-based virtual screening tools can identify potential binding proteins but suffer from low hit rates due to neglect of protein conformational flexibility and real-world experimental conditions^[26]^. More recently, AI-driven denovo protein design has generated substantial excitement, with platforms such as RFdiffusion^[27]^ and ProteinMPNN^[28]^ enabling rapid creation of novel binders. However, these methods predominantly target protein-protein interactions, and their application to small-molecule recognition remains limited by the challenge of designing complementary binding pockets for low molecular-weight, often hydrophobic ligands. In contrast, natural pocket proteins have been pre-optimized by evolution specifically for small-molecule binding, offering practical advantages in terms of stability, solubility, and manufacturing readiness. In addition, no integrated platform currently links rapid recognition-element discovery for small molecules with direct translation into portable POCT sensing.

Here, we present DWCLASS, a strategy that connects AI-guided identification and repurposing of natural pocket proteins with portable MetaSPR sensing (Fig. 1). The workflow combines multimodal affinity prediction across approximately 16,000 protein structures, molecular simulations, and eight-channel experimental screening. We examine thiamine (vitamin B1, VB1) and 25-hydroxyvitamin D3 [25(OH)D3] as chemically distinct model analytes^[29][30]^, selecting 1GAI and 1ADD for competitive detection. Seven of the 16 tested analyte–protein pairs produced binding responses, corresponding to a 43.8% selected-set hit fraction. Incorporating the selected proteins into an AuNP-amplified, acoustofluidics-assisted device enabled two-analyte measurements with a 240 s acquisition step and smartphone readout. Serum measurements provide an application of this workflow, connecting protein-library screening to a functional portable assay.

**Fig. 1.**
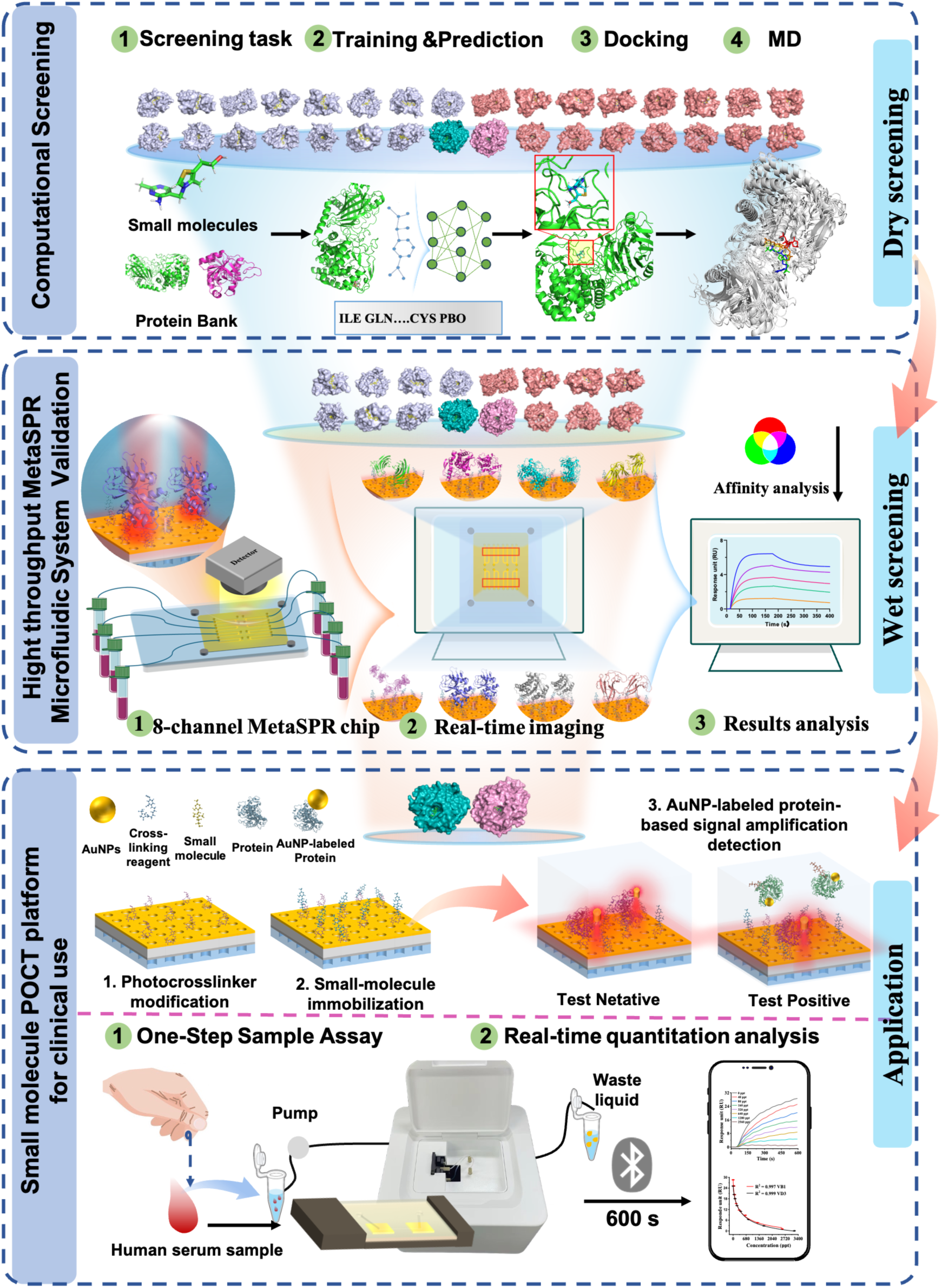
High-Throughput DWCLASS Strategy for Efficient Small-Molecule Pocket Protein Screening and Ultrasensitive Small-Molecule Detection: High-efficiency small-molecule pocket protein screening based on the PLCA model and molecular dynamics simulations; Rapid affinity evaluation of candidate pocket proteins using an 8-channel MetaSPR microfluidic molecular interaction analysis system; Ultrasensitive and rapid quantitative detection of small-molecules achieved by integrating high-affinity pocket proteins with an acoustofluidics-assisted MetaSPR sensing POCT system.

## 2. Results and Discussion

### 2.1. The PLCA Model: AI-enabled reverse screening identifies natural pocket proteins for target small molecules

Traditional protein-ligand affinity prediction models are designed for the task of identifying small molecules that bind to a given protein. They are optimized to rank small molecules against a single protein target, not to rank proteins against a single small molecule. This fundamental mismatch leads to poor performance when applied to pocket protein screening. To address this, we developed the PLCA model, a multimodal deep learning architecture specifically trained for the reverse task of small-molecule-to-protein binding affinity prediction.

The PLCA model integrates three complementary representations of protein-ligand complexes. As illustrated in **Fig. 2a**: (1) Multi-Perspective Data Representation, which encodes amino acid sequences and 3D protein structural topologies into compatible feature spaces; (2) Encoder Module, which leverages deep learning to mine latent interaction patterns from the fused features; and (3) Readout Module, which maps the encoded features to quantitative binding affinity values. This design rationale aligns with the biological reality that protein-ligand binding is governed by both sequence-derived biochemical properties (e.g., hydrophobicity, charge distribution) and structure-dependent spatial constraints (e.g., binding pocket geometry, intermolecular hydrogen bonding).

**Fig. 2.**
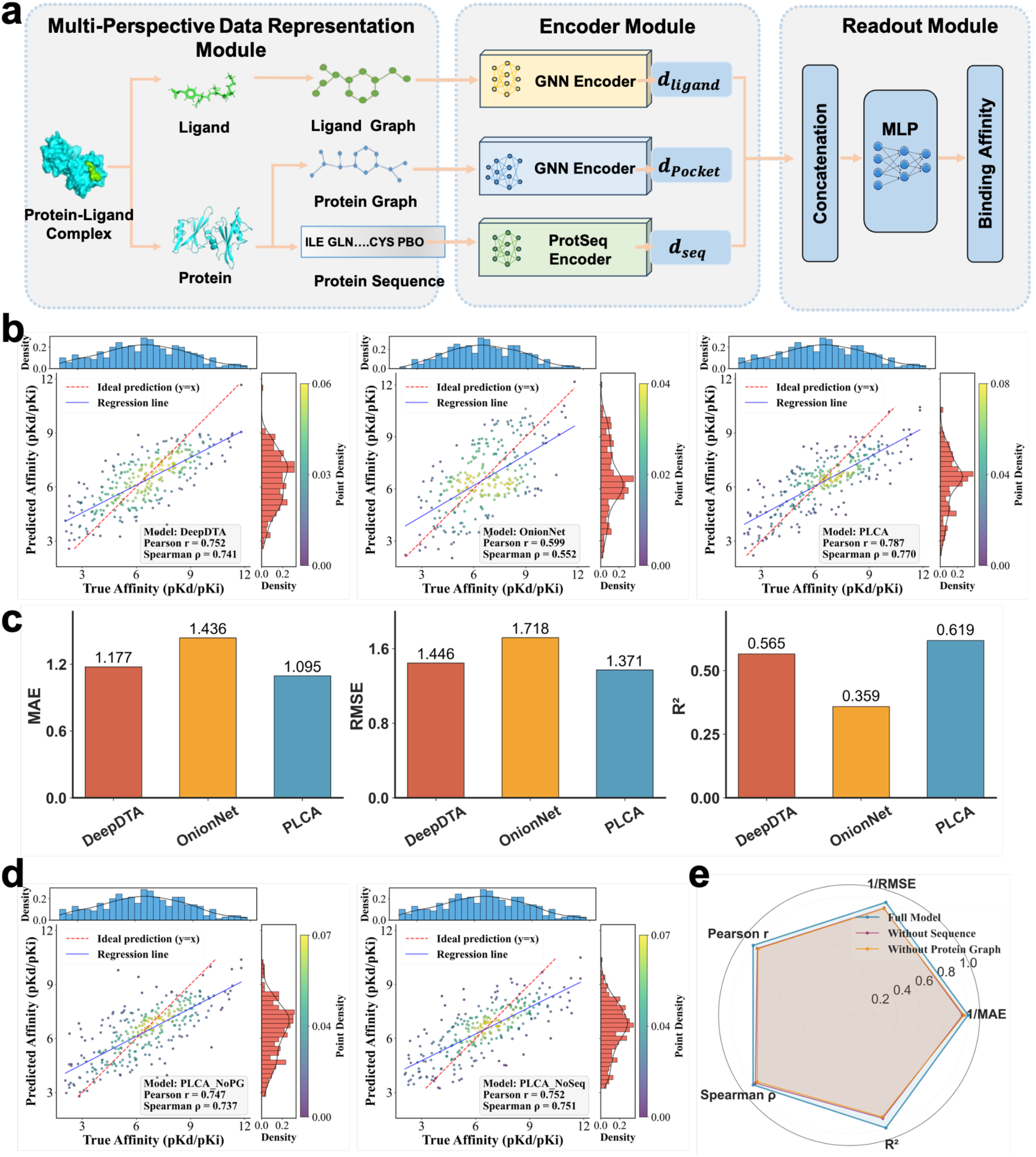
Design and evaluation of the PLCA model. (a) PLCA framework for high-throughput screening of small-molecule pocket proteins, built on a multi-modal architecture that integrates structural geometry with biochemical sequence information. (b) Correlation analysis of predicted versus experimental binding affinities for DeepDTA, OnionNet and PLCA on the PDBbind-2016 test set. (c) Comparative performance of the three models in terms of MAE, RMSE and R². (d) Correlation results of PLCA when sequence input or pocket graph input is ablated. (e) Ablation experimental outcomes comparing the full PLCA model with its variants lacking sequence information or pocket graph information.

PLCA was evaluated alongside DeepDTA^[31]^, a sequence-based model, and OnionNet^[32]^, a structure-based model, on the PDBbind-2016 dataset^[33]^. Figure 2b compares predicted and experimental affinity values, and Figure 2c summarizes the corresponding prediction errors and R² values.

PLCA predictions clustered more closely around the reference diagonal than those of the two comparison models (Fig. 2b). On this test set, PLCA achieved MAE = 1.095, RMSE = 1.371, and R² = 0.619, compared with MAE = 1.177, RMSE = 1.446, and R² = 0.565 for DeepDTA, and MAE = 1.436, RMSE = 1.718, and R² = 0.359 for OnionNet (Fig. 2c). These results support combining sequence and local structural information for affinity prediction within the evaluated benchmark.

To dissect the individual contributions of the sequence and structural components to PLCA’s performance, we conducted systematic ablation experiments. Specifically, we generated two ablated variants of the full model: one lacking the Sequence Input (PLCA-NoSeq) and another lacking the Pocket Graph Input (PLCA-NoPG). The performance of these variants was compared to the full PLCA architecture under the same experimental conditions, with results presented in **Fig. 2d, e**.

Removing the sequence input increased MAE from 1.0954 to 1.1464 and reduced R² from 0.6186 to 0.5659. Pearson and Spearman correlations decreased from 0.7865 to 0.7522 and from 0.7698 to 0.7511, respectively (Fig. 2d,e). The sequence representation thus contributed information complementary to the structural inputs in this benchmark.

Removing the pocket-graph input increased RMSE from 1.3708 to 1.4520 and residual standard deviation from 1.3587 to 1.4470 (Fig. 2d,e). Local pocket geometry therefore contributed to the accuracy of the combined representation. Together, the two ablations support retaining both sequence and pocket-graph inputs in the subsequent screening workflow.

We next applied PLCA to rank approximately 16,000 candidate protein structures for VB1 and 25(OH)D3. Model scoring was completed in less than 30 min, and the top 40 candidates for each analyte were selected for further computational evaluation (Table S3). This library-scale ranking provided a tractable starting point for docking, molecular simulations, and experimental binding measurements.

### 2.2. Closing the Dry-Wet Gap: From Computational Predictions to Experimental Hits

A major limitation of traditional AI virtual screening is the low hit rate when predictions are validated experimentally. This is due to several factors: static crystal structures do not capture protein conformational flexibility, models ignore experimental conditions such as pH and temperature, and there is no feedback loop between computational predictions and experimental results. To address this, we designed a closed-loop dry-wet validation pipeline that combines MD simulations with high-throughput MetaSPR analysis, significantly improving hit rates and reducing experimental waste.

Starting from the top 40 PLCA-ranked candidates for thiamine and 25(OH)D₃, we first removed proteins with >90% sequence homology to avoid structural redundancy, resulting in 16 non-homologous candidates per analyte (**Fig. S1 and S2**). We then performed molecular docking using AutoDock Vina to confirm static binding feasibility. All 32 candidates showed docking scores < 0 kcal·mol⁻¹, indicating spontaneous binding, with 31 candidates scoring < -5 kcal·mol⁻¹ (Supplementary **Tables S4**). And the docking results are shown in **Fig. 3a,b**. Next, we conducted 100 ns MD simulations using GROMACS to evaluate the dynamic stability of the protein-ligand complexes. We quantified binding free energy using the MMPBSA method, as well as hydrogen bond persistence and protein-ligand contact area over the simulation trajectory(**Fig. 3c, d**, Supplementary **Tables S5**). All candidates maintained stable complexes (no small molecule dissociation), with a subset standing out for balanced affinity and stability: for thiamine, 1J5I had the lowest binding free energy (-35.49 kcal·mol⁻¹), paired with 1.35 average HBs and a ∼73 Å² contact area; 1GAI (Glucoamylase-471)^[34]^ followed (-24.89 kcal·mol⁻¹, 3.2 HBs, ∼160 Å²); and 2ADA had the heighest contact area with 162 Å². For 25(OH)D₃, 4CSD showed the most favorable free energy (-48.09 kcal·mol⁻¹) and ∼132 Å² contact area,. 3D Free energy landscape (3D_FEL) diagrams (from RMSD/Rg) and conformational snapshots (0/25/50/75/100 ns) further validated stability: top thiamine candidates and 25(OH)D₃ candidates formed single, compact low-energy clusters (dark purple/blue regions) with no binding position shifts (**Fig. 3e, f**). The corresponding 2D projections of the free energy landscapes are provided in Supplementary **Figs. S3** and **S4**. This step eliminated candidates that showed unstable binding or large conformational changes, narrowing the pool to 8 high-priority candidates per analyte.

**Fig. 3.**
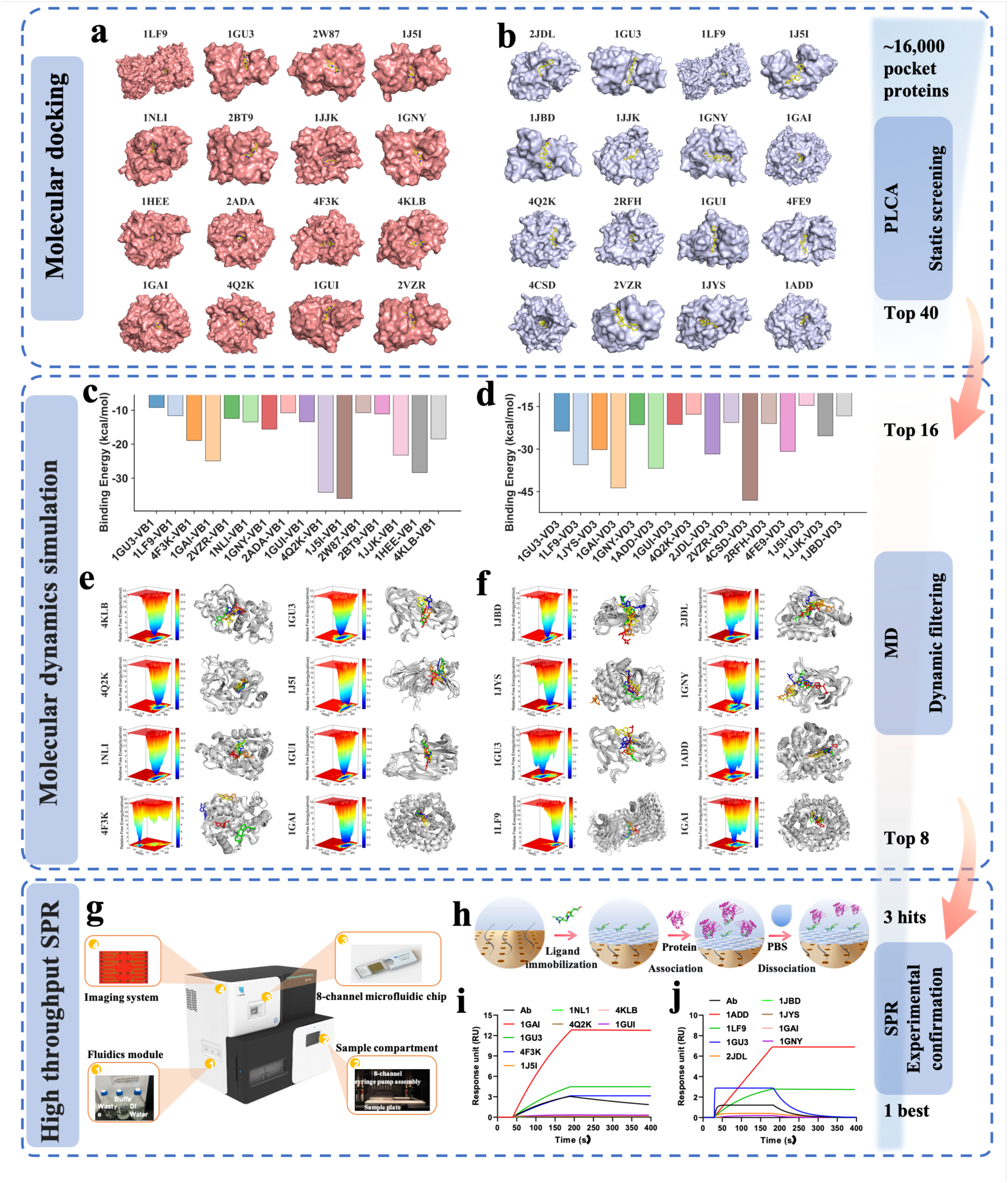
Affinity Prediction of Protein-ligand via Molecular Docking and Molecular Dynamics (MD) Simulations. (a) Molecular docking conformations of thiamine with 16 candidate pocket proteins. (b) Molecular docking conformations of 25(OH)D₃ with 16 candidate pocket proteins. (c) Results of molecular dynamics (MD) simulations for thiamine and its 16 candidate pocket proteins of MMPBSA-calculated binding free energy. (d) Results of molecular dynamics (MD) simulations for 25(OH)D₃ and its 16 candidate pocket proteins of MMPBSA-calculated binding free energy. (e) Free energy landscapes (FELs) and conformational changes at 0, 25, 50, 75, and 100 ns from MD simulations of thiamine with 16 candidate pocket proteins. (f) Free energy landscapes (FELs, 3D) and conformational changes at 0, 25, 50, 75, and 100 ns from MD simulations of 25(OH)D₃ with 16 candidate pocket proteins. (g) High-throughput MetaSPR imaging detection system; (h) Small Molecule Pocket Protein Affinity Detection Schematic; (i) Rapid affinity comparison analysis results comparison of thiamine with 8 candidate proteins (all at 300 nM protein concentration); (j) Rapid affinity comparison analysis results comparison of 25(OH)D₃ with 8 candidate proteins (all at 300 nM protein concentration).

We then validated these 8 candidates experimentally using an 8-channel MetaSPR microfluidic system, which allows parallel, label-free measurement of binding kinetics in real time(**Fig. 3gh**). For thiamine, 3 out of 8 candidates showed specific binding, resulting in a hit rate of 37.5%(**Fig. 3i and Fig.S5**). For 25(OH)D₃, 4 out of 8 candidates showed specific binding, resulting in a hit rate of 50.0%(**Fig. 3j and Fig.S6**). The overall hit rate across both analytes was 43.8%, a 2.5-fold improvement over the 17.5% hit rate reported for the state-of-the-art DrugCLIP model. This dramatic improvement demonstrates the power of the closed-loop dry-wet pipeline: by combining AI screening with MD simulation and high-throughput experimental validation, we are able to efficiently filter out false positives and focus experimental efforts on the most promising candidates.

### 2.3. Binding characterization and interaction analysis of repurposed pocket proteins

We next characterized the responsive candidates to select proteins for assay development. The selected scaffolds were derived from glucoamylase (1GAI)^[34]^ and adenosine deaminase (1ADD)^[35]^, which were repurposed here as recognition reagents for VB1 and 25(OH)D3, respectively. In the MetaSPR surface assay, 1GAI generated the largest response among the VB1 candidates, approximately fourfold that of the commercial VB1 antibody included as a reference. For 25(OH)D3, 1ADD produced the largest response, approximately sevenfold that of the tested commercial antibody (Fig. 4a,b). These measurements identified 1GAI and 1ADD as candidates for subsequent sensing experiments.

**Fig. 4.**
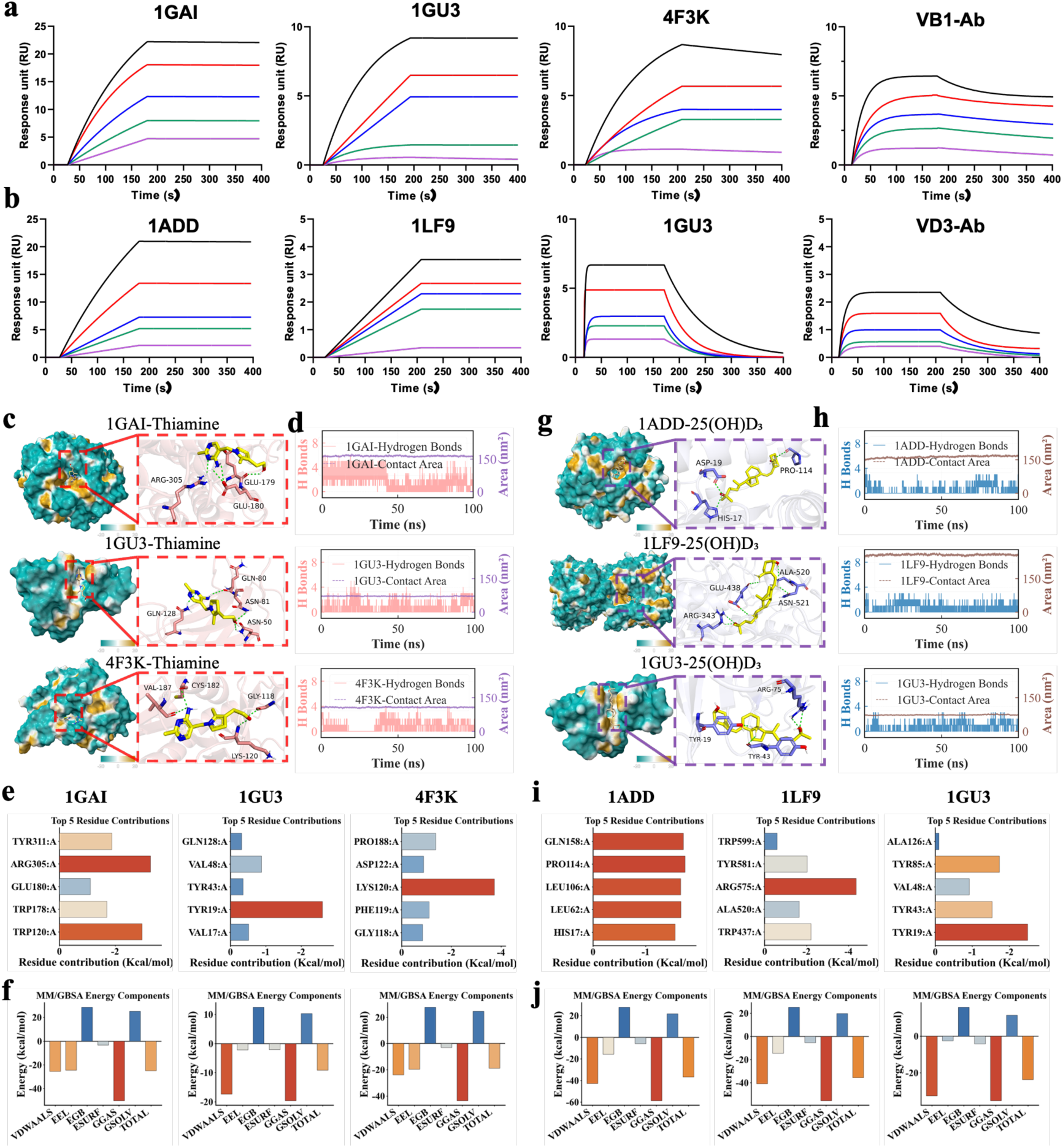
High-Throughput MetaSPR Affinity Validation and Binding Mechanism Analysis of Pocket Proteins and small molecules. (a) MetaSPR affinity analysis results of thiamine with 3 potential binding pocket proteins (1GAI, 1GU3, and 4F3K) and commercial thiamine-Ab at different concentrations; (b) MetaSPR affinity analysis results of 25(OH)D₃ with 3 potential binding pocket proteins (1ADD, 1LF9, and 1GU3) and commercial 25(OH)D₃-Ab at different concentrations; (c) Molecular docking conformation analysis of thiamine with 3 potential binding pocket proteins (1GAI, 1GU3, 4F3K); (d) Hydrogen bond and contact area changes of thiamine with 3 potential binding pocket proteins (1GAI, 1GU3, 4F3K) during molecular dynamics simulations; (e) Amino acid residue energy contribution of thiamine with 3 potential binding pocket proteins (1GAI, 1GU3, 4F3K); (f) Average binding free energy of thiamine with 3 potential binding pocket proteins (1GAI, 1GU3, 4F3K), where VDWAALS, EEL, EGB, ESURF, GGAS, GSOLV, and TOTAL represent van der Waals force, electrostatic energy, polar solvation energy, non-polar solvation energy, molecular mechanics term, solvation energy term, and total average binding free energy, respectively;(g) Molecular docking conformation analysis of 25(OH)D₃ with 3 potential binding pocket proteins (1ADD, 1LF9, 1GU3); (h) Hydrogen bond and contact area changes of 25(OH)D₃ with 3 potential binding pocket proteins (1ADD, 1LF9, 1GU3) during molecular dynamics simulations; (i) Amino acid residue energy contribution of 25(OH)D₃ with 3 potential binding pocket proteins (1ADD, 1LF9, 1GU3); (j) Average binding free energy of 25(OH)D₃ with 3 potential binding pocket proteins (1ADD, 1LF9, 1GU3), where VDWAALS, EEL, EGB, ESURF, GGAS, GSOLV, and TOTAL represent van der Waals force, electrostatic energy, polar solvation energy, non-polar solvation energy, molecular mechanics term, solvation energy term, and total average binding free energy, respectively.

To examine possible molecular interactions underlying the binding responses, we combined docking, 100 ns MD simulations, and per-residue energy decomposition. In the modeled 1GAI–VB1 complex, contacts involving GLU179, GLU180, and ARG305 were associated with an average of 3.2 hydrogen bonds (Fig. 4c,d). The 1GU3 carbohydrate-binding module^[36]^ and 4F3K showed different contact patterns. Per-residue analysis identified five residues with contributions more favorable than −1 kcal mol−1 for 1GAI, compared with one for 1GU3 and three for 4F3K (Fig. 4e). The corresponding endpoint energy components provide a comparative description of the modeled complexes (Fig. 4f).

For 25(OH)D₃-binding proteins, 1ADD formed a stable complex with an average of 3 HBs to residues HIS-17, ASP-19, and PRO-114, with an average contact area of ∼195 Å² (**Fig. 4g, h**). Balanced contributions from hydrogen bonding and hydrophobic interactions underpinned its exceptional affinity. Notably, 1LF9 formed an average of 6 HBs, the highest among all candidates, but suffered from severe steric hindrance caused by its deeper, narrower binding pocket that restricted 25(OH)D₃ access, resulting in a lower overall binding free energy (-35.50 kcal/mol) than 1ADD (-36.78 kcal/mol). Per-residue energy decomposition analysis (**Fig. 4i**) revealed that 1ADD had 14 residues contributing to binding, the most of any 25(OH)D₃ candidate, explaining its superior performance (**Fig. 4j**).

A particularly interesting finding was the dual specificity of 1GU3, which bound both thiamine and 25(OH)D₃ with high affinity. Structural analysis revealed that 1GU3 contains two distinct, non-overlapping binding pockets: a hydrophilic pocket for thiamine and a hydrophobic pocket for 25(OH)D₃, suggesting either multiple accessible pocket environments or reduced selectivity relative to 1GAI and 1ADD. Therefore, it was not selected as the final recognition element for multiplexed sensing.

In both thiamine and 25(OH)D₃ complexes, the binding pocket completely encloses the small molecule, resulting in shape complementarity. This near-perfect shape complementarity, refined over hundreds of millions of years of evolution, is the primary structural basis for the exceptionally high affinity of natural pocket proteins. Unlike antibodies, which evolve through somatic hypermutation over weeks to months with selection pressure limited to binding affinity, natural pocket proteins have evolved under strong selective pressure for not only high affinity and specificity but also structural stability and efficient folding.

Complementary data confirmed the exceptional stability of these protein-ligand complexes: RMSD values remained < 2 Å for all active complexes over the entire 100 ns MD simulation, and solvent-accessible surface area (SASA) analysis showed that > 90% of the small molecule surface is buried upon binding (Supplementary **Figs. S7–S22**). Specificity validation confirmed that 1GAI exclusively binds thiamine and 1ADD exclusively binds 25(OH)D₃, with no detectable cross-reactivity with structurally similar molecules (Supplementary **Fig. S23**), a critical requirement for multiplexed detection.

These results have important implications for biosensing. Antibodies are the gold standard for recognition elements in immunoassays, but they are limited by the constraints of the immune system. Natural pocket proteins, on the other hand, have evolved over millions of years to bind small molecules with high affinity and specificity. By using AI to discover these proteins, we can tap into a vast, untapped reservoir of recognition elements that outperform antibodies for many small-molecule targets. This eliminates the need for immunization and reduces development time.

### 2.4. Breaking the Sensitivity-Speed-Portability Trade-off with an Acoustofluidics-Enhanced MetaSPR POCT Platform

Armed with these high-affinity natural pocket proteins, we set out to address the most intractable challenge in small-molecule biosensing: the **sensitivity-speed-portability** trade-off that has defined POCT technology for decades. For small molecules, this trade-off is particularly severe: high-sensitivity detection requires bulky laboratory instruments with assay times of hours; rapid assays rely on antibodies that lack sufficient affinity; and portable devices sacrifice performance for simplicity.

We chose metasurface plasmon resonance (MetaSPR) as the detection technology because it is the only platform that can potentially reconcile these three conflicting requirements. Unlike traditional SPR, which relies on bulky prism coupling, MetaSPR uses engineered nanostructures to generate localized surface plasmon resonance, enabling label-free, real-time detection with high sensitivity in a miniaturized format. However, conventional MetaSPR sensors suffer from two fundamental limitations that prevent their application to trace small-molecule detection: (1) slow binding kinetics due to diffusion-limited mass transport, and (2) low signal intensity for low-molecular-weight analytes. To overcome these limitations, we developed a synergistic platform that integrates three core innovations: a novel indirect competitive assay tailored for pocket proteins, gold nanoparticle (AuNP) signal amplification, and sharp-edge induced acoustic streaming mixing.

The MetaSPR chip was fabricated via a replica-molding nanoimprint process combined with noble metal deposition, enabling cost-effective and scalable wafer-level production critical for practical POCT applications. Titanium was used as an adhesion layer to enhance golddeposition stability, while gold was selected for its minimal non-specific binding, strong plasmonic effect, and long-term operational stability. SEM characterization confirmed uniform nanostructural parameters across the chip surface, ensuring excellent reproducibility (**Fig. 5a**). FDTD simulations show negligible electric field enhancement at 550 nm (indicating low sensitivity to refractive index (RI) changes) and significant field localization at 640 nm (**Fig. 5b**), validating 640 nm as the optimal detection wavelength for maximizing signal response to molecular binding events. We tested solutions with different sucrose concentrations (**Fig. 5d, e**). The differential OD spectra showed a distinct, concentration-dependent trend, with the maximum signal difference at 640 nm, consistent with FDTD simulation results. The chip achieved a linear response to sucrose concentration (R² > 0.99), with a detection limit (LOD) of 0.03%. This ultrahigh RI sensitivity lays the foundation for detecting trace small molecules, as their binding to pocket proteins induces subtle surface RI shifts that the MetaSPR chip can resolve.

**Fig. 5.**
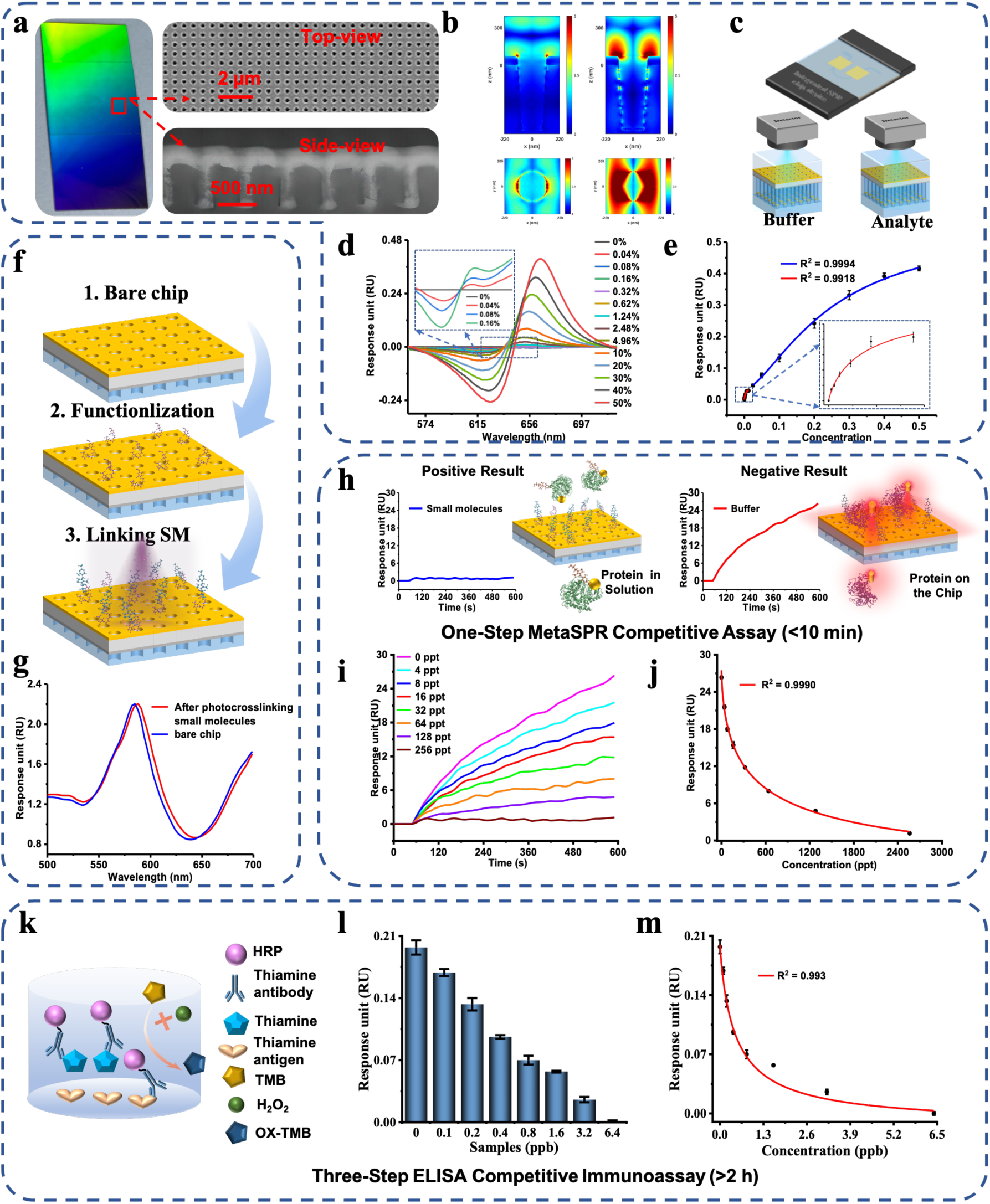
Development of a Small-Molecule Detection Sensor Based on Nanocup Array MetaSPR. (a) Optical image and SEM characterization (top and side views) of the nanocup array MetaSPR sensor chip; (b) Comparison of electric field distributions calculated by FDTD at non-responsive and maximum wavelength response positions; (c) Integrated microfluidic MetaSPR sensing device and detection schematic; (d) OD spectral curves of the MetaSPR chip in response to different sucrose concentrations; (e) Fitting curve of the MetaSPR chip’s response (maximum minus minimum OD value) to different sucrose concentrations; (f) Schematic of small-molecule modification via photo-crosslinking; (g) OD spectral changes before and after modification; (h) Schematic of the positive and negative sample detection using the MetaSPR sensor chip; (i) Real-time binding responses of the pocket protein biosensor toward a concentration gradient of thiamine in buffer. (j) Fitting curve results of the MetaSPR sensor chip for thiamine standard samples; (k) Principle of sharp-edge induced acoustic streaming: PZT excitation causes the sharp tips to oscillate, generating steady micro-vortices that convect targets toward the sensing interface; (l) Response curves of small molecule detection with different acoustofluidics frequencies(0 Khz, 4 KHz, 8 KHz, 12 KHz); (m) Comparison of signal responses under different acoustofluidics frequencies after 200 s of reaction; (n) Comparison of time consumption under different acoustofluidics frequencies when the response signal reaches 20 RU; (o) Protocol for thiamine quantification via ELISA; (p) Response results of ELISA for thiamine standard samples; (q) Fitting curve results of ELISA for thiamine standard samples.

Small molecules were immobilized on the MetaSPR surface using the photo-crosslinking procedure (Fig. 5f). The spectral shift after modification was consistent with a change in the sensing interface (Fig. 5g). This functionalized surface provides the capture ligand for competition between soluble analyte and surface-presented analyte

The competitive assay converts small-molecule recognition into capture of AuNP-labeled protein (Fig. 5h; Fig. S24). Labeled pocket proteins were mixed with the sample before exposure to the functionalized surface. Soluble analyte occupies recognition sites and reduces subsequent capture by surface-presented analyte, generating a decreasing response as sample concentration increases. A reference channel lacking immobilized analyte was used to reduce shared background and nonspecific signals. The resulting format couples the recognition properties of the selected proteins with an amplified optical readout.

For thiamine detection, distinct concentration-dependent signal gradients were observed within 600 s (**Fig. 5i**), demonstrating rapid analytical capability. Using the average of the last five response signals from 600 s kinetic curves, a 4-parameter logistic (4-PL) fitting curve was constructed with a high correlation coefficient (R² = 0.9990) (**Fig. 5j, Fig. S25)**. The LOD for thiamine was 0.72 ppt calculated by interpolating the blank sample signal plus three times its standard deviation (SD) into the fitting curve.

A sharp-edge acoustic mixer driven by a piezoelectric transducer was integrated into the microchannel (Fig. 5k; Fig. S26). The simulations predicted local streaming velocities of 10–20 mm/s, compared with a bulk velocity of approximately 2.2 mm/s. Among the tested actuation frequencies, 8 kHz produced the largest response: the signal increased 3.38-fold, and the time to reach 20 RU decreased by 54% relative to the unactuated condition (Fig. 5l–n). Acoustic mixing thus improved the response rate within the tested device. The subsequent portable measurements used a 240 s acquisition interval.

This sensitivity is 6260-fold higher than a recently reported antibody-based ELISA (LOD = 4.51 ng/mL)^[10]^ and 75-fold higher than traditional ELISA (LOD = 54.51 ppt) (**Fig. 5o, q**). Additionally, the assay time (10 min) was 12-fold shorter than that of ELISA (∼120 min), highlighting the sensor’s superior speed and sensitivity through the integration of DWCLASS pocket proteins (high affinity/specificity) and MetaSPR technology (rapid signal transduction).

Selectivity, stability, and reproducibility were further validated to confirm practical applicability. Challenging the sensor with structurally similar interfering molecules yielded no significant signal changes (**Fig. S27-S29**), confirming high selectivity attributed to the pocket proteins’ specific binding (validated in Section 3.3). After 6 months of storage at 25°C, the sensor maintained consistent signal responses to thiamine (**Fig. S30**), indicating storage stability under these conditions. Evaluations across chip regions and fabrication batches showed low coefficients of variation in signal responses (Figs. S31–S33), supporting the uniformity of the fabricated chips. Thiamine calibration curves obtained using chips from three fabrication batches also exhibited similar concentration-dependent response profiles (Fig. S34). For quantitative sample analysis, we nevertheless recommend establishing an independent calibration curve for each fabrication batch to account for potential batch-dependent variation in sensor response.

### 2.5. Clinical Validation in Human Serum Samples

We integrated the assay components into a portable two-analyte device for serum measurements. The device combines the selected proteins, 1GAI and 1ADD, with spatially separated MetaSPR sensing regions, a microfluidic cartridge, acoustic mixing, and smartphone readout (Fig. 6a–g). This configuration extends the workflow from candidate selection to an assembled analytical device.

**Fig. 6.**
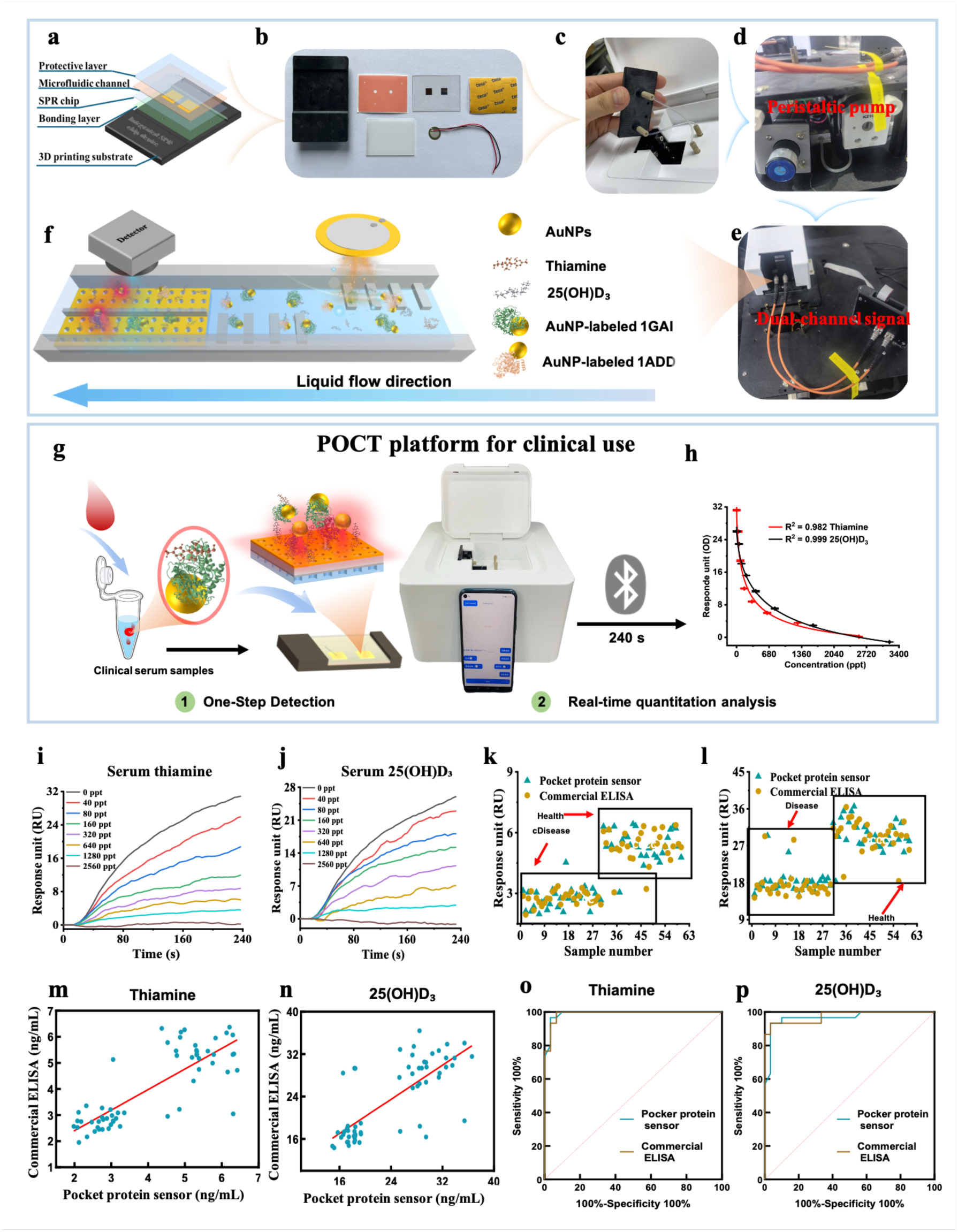
Development of an acoustofluidics-Enhanced, Ultrasensitive Multiplexed Small-Molecule Detection Device and Its Clinical Application. (a) Schematic design of the integrated microfluidic MetaSPR detection cartridge; (b) Physical image of components for the acoustofluidics-integrated microfluidic MetaSPR detection cartridge; (c) Photograph of the rapid cartridge loading module of the POCT device; (d) Miniaturized peristaltic pump module for microfluidic flow control; (e) Dual-optical-path detection module with reference and measurement channels; (f) Enlarged schematic of the sharp-edge array microchannel with spatially partitioned dual-analyte detection zones; (g) Schematic of the one-step small-molecule POCT platform based on screened pocket proteins and MetaSPR; (h) Standard fitting curves of detection results for thiamine and 25(OH)D₃ standards in serum.(i-j) Real-time binding curves for different concentrations of thiamine (i) and 25(OH)D₃ (j) in human serum; (k) Comparison of thiamine detection results between ELISA and the pocket protein-based POCT device in human serum samples (n = 60); (l) Comparison of 25(OH)D₃ detection results between ELISA and the pocket protein-based POCT device in human serum samples (n = 60); (m) Correlation between the pocket protein sensor developed in this study and commercial ELISA for measuring thiamine levels; (n) Correlation between the pocket protein sensor developed in this study and commercial ELISA for measuring 25(OH)D₃ levels; (o) ROC curves for thiamine detection by ELISA and the pocket protein-based POCT device; (p) ROC curves for 25(OH)D₃ detection by ELISA and the pocket protein-based POCT device.

The device features a modular, user-centric design optimized for point-of-care operation. The core component is a disposable microfluidic detection cartridge fabricated via low-cost 3D printing and lamination (**Fig. 6a, b**). This single-use cartridge eliminates cross-contamination between samples, requires only 100 μL of serum per test, and can be loaded into the device in less than 10 seconds (**Fig. 6c**). A miniaturized peristaltic pump provides precise control of sample flow rate (40 μL/min), eliminating the need for external pressure sources (**Fig. 6d**). To ensure stable performance in variable environmental conditions, we integrated a dual-optical-path system with a reference channel that corrects for baseline drift caused by temperature fluctuations and non-specific adsorption (**Fig. 6e**).

To enable simultaneous detection of multiple biomarkers from a single sample—a critical requirement for clinical diagnosis where sample volume is often limited—the MetaSPR chip’s detection zone is spatially partitioned into two independent sub-regions. thiamine and 25(OH)D₃ were immobilized in separate sub-regions via the photo-crosslinking method established in Section 2.4, allowing parallel quantification of both analytes from a single sample injection (**Fig. 6f**). This design eliminates the need for multiple chip runs or sample splitting, reducing assay time and sample consumption by 50% compared to single-analyte tests.

Binding curve analysis confirmed a clear inverse relationship between signal intensity and analyte concentration for both thiamine and 25(OH)D₃, with 4-parameter logistic (4-PL) fitting curves showing high correlation coefficients (R² = 0.997 for thiamine, R² = 0.999 for 25(OH)D₃; **Fig. S35-S37**). The device achieved LODs of 1.76 ppt (thiamine) and 1.41 ppt (25(OH)D₃) in buffer. These values are consistent with the single-analyte sensor’s performance (thiamine: 0.72 ppt) and 12.33-fold more sensitive than the gold standard LC-MS/MS (25(OH)D₃: 17.38 ng/mL) ^[7]^. This sensitivity improvement stems from the synergistic effect of the DWCLASS strategy: the AI-screened pocket proteins (1GAI/1ADD) provide high-affinity recognition, while MetaSPR’s plasmonic enhancement and acoustofluidics-assisted mixing amplify the signal of trace small molecules.

To validate performance in complex clinical matrices, we optimized the assay buffer to mitigate matrix interference from serum proteins, lipids, and metabolites. The device maintained distinct concentration-dependent signal gradients for both thiamine and 25(OH)D₃ in human serum (**Fig. 6g, h** and **Fig. S38**), with LODs of 2.98 ppt for thiamine and 3.82 ppt for 25(OH)D₃. This represents only a 1.7-fold and 2.7-fold increase in LOD compared to buffer, respectively—far superior to the 10-100-fold increase typically observed for antibody-based immunoassays. Real-time dynamic curves showed clear differentiation between clinically relevant concentrations within 240 seconds (**Fig. 6i, j**), confirming that serum components do not significantly interfere with pocket protein-small molecule binding. This exceptional robustness is attributed to the high specificity of the AI-discovered pocket proteins and the dual-optical-path system’s ability to suppress non-specific adsorption.

Serum samples were measured using the pocket-protein device and commercial ELISA (Fig. 6k,l; Table S6). Passing-Bablok regression analysis further confirmed strong agreement between the two platforms (R² > 0.99 for both analytes; **Fig. 6mn**), validating that the device’s results are clinically actionable. Receiver Operating Characteristic (ROC) curve analysis showed exceptional diagnostic performance (**Fig. 6op**): for thiamine, the device achieved 96.67% sensitivity and 96.67% specificity (AUC = 0.99), matching ELISA’s sensitivity (100.00%) while showing improved specificity (ELISA: 93.33%); for 25(OH)D₃, it reached 93.33% sensitivity and 96.67% specificity (AUC = 0.97), outperforming ELISA’s specificity (93.33%).

The portable measurements complete the connection between computational protein selection and assay operation. The selected natural scaffolds provide the recognition reagents, the functionalized MetaSPR surface supports competitive capture, and AuNP labeling and acoustic mixing link that recognition to an optical signal. The integrated acoustofluidics system overcomes the classic speed-sensitivity trade-off, enabling rapid assays without compromising accuracy. It thus overcomes the key limitations of traditional methods and offers a scalable, cost-effective alternative for high-quality small-molecule detection across diverse settings, demonstrating an approach to monitoring trace biomarkers in health and disease.

## 3. Conclusion

This work establishes DWCLASS as an integrated strategy for identifying and repurposing natural pocket proteins for small-molecule biosensing. Through the demonstrations with VB1 and 25(OH)D3, AI-guided candidate identification and experimental selection were connected to the construction of competitive MetaSPR assays and portable serum measurements. These findings demonstrate that selected natural protein scaffolds can serve new analytical functions when their recognition properties are incorporated into an appropriate sensing interface.

The significance of this approach lies in connecting the structural diversity of natural proteins with the requirements of analytical device design. Target-directed searching provides access to candidate recognition elements within existing protein libraries, while experimental evaluation establishes their suitability for assay development. The resulting analytical performance depends on both molecular recognition and its integration with surface presentation, signal amplification, and transport control. Developing these functions together provides a route for translating computationally prioritized interactions into measurable signals and functional sensing devices.

The present demonstration is limited to two model analytes, and its applicability across a broader range of chemical structures remains to be established. Recognition selectivity and performance in biological samples will also depend on the chosen scaffold and assay conditions. Further work should therefore expand analyte coverage, examine the structural determinants of recognition, and combine protein optimization with interface engineering. Integration of sample preparation and evaluation in larger, independent sample cohorts will help define the practical scope of the approach and support its development for specific analytical applications.

DWCLASS provides a complementary route to recognition-reagent development through natural-protein repurposing. Its broader potential lies in connecting existing protein-structure resources with the design of analytical interfaces. This connection offers opportunities to expand the repertoire of protein-based recognition elements and develop portable small-molecule biosensors whose molecular components and device architectures are designed in concert.

## 4. Materials and methods

### 4.1. Prediction Model Design for Protein-Ligand Complex Binding Affinity

The **PLCA** model is designed to predict the **P**rotein-**L**igand **C**omplexes binding **A**ffinity which integrates structural and sequential information. The architecture consists of three primary stages: **Multi-Perspective Data Representation**, **Encoder Module**, and **Readout Module**.

**① Multi-Perspective Data Representation:** To capture the comprehensive features of the protein-ligand interaction, we represent the input complex from three distinct perspectives:

- **Ligand Graph *G_L_*:** The ligand is represented as a molecular graph *G_L_*=(*V_L_*,*E_L_*), where *V_L_* denotes the set of atoms (nodes) and *E_L_* denotes the chemical bonds (edges). Each node *i*∈*V_L_*is initialized with a feature vector *x_i_* encoding atomic properties.
- **Pocket Graph *G_P_*:** Rather than using the entire protein, we focus on the binding pocket to reduce noise. The pocket is modeled as a graph ***G_P_*=**(*V_P_*,*E_P_*): where nodes *V_P_* represent amino acids or heavy atoms within a specific distance from the ligand, and edges *E_P_* are defined based on spatial proximity.
- **Protein Sequence *S*:** To preserve the primary structure information, the protein is also represented as a sequence of amino acid residues, denoted as *S*={*s*_1_,*s*_2_,…,*s_n_*}.

**②Encoder Module:** This module extracts high-level latent embeddings from the raw graph and sequence representations.

- **Graph Neural Network (GNN) Encoders:** We employ two separate GNN encoders for the ligand and the pocket graphs. For a node *i* in a graph, the message-passing and update rule at layer *l*+1 is defined as:

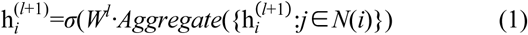

where *N*(*i*) represents the neighbors of node *i*, *W^l^* is a learnable weight matrix, and *σ* is a non-linear activation function (e.g., ReLU). After *l* layers of graph convolutions, a global pooling (Readout) operation is applied to obtain the fixed-dimension graph embeddings:

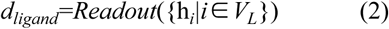

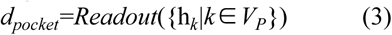
- **Protein Sequence Encoder:** The protein sequence ***S*** is processed through a sequence encoder (e.g., Transformer, CNN, or BiLSTM) to capture the biochemical context:

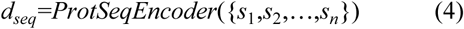

where *d_seq_* is the protein sequence embedding vector.

**③Readout Module**: The final stage fuses the multi-perspective embeddings to perform the binding affinity prediction task of protein-ligand complexes.

- **Feature Concatenation:** The three embedding vectors (*d_ligand_*, *d_pocket_*, and *d_seq_*) are concatenated into a single joint representation vector *z*:

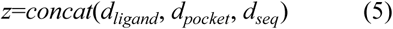

where *concat*(·) denotes the concatenation operation.

**Affinity Prediction**: The concatenated vector *z* is passed through a Multi-Layer Perceptron (MLP) consisting of several fully connected layers with dropout and activation functions:

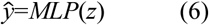

where *ŷ* is the predicted binding affinity value (e.g., *pK_d_*,*pK_i_*).

**Training Objective:**The model is trained end-to-end by minimizing the Mean Squared Error (MSE) between the predicted affinity *ŷ* and the experimental ground truth *y*:

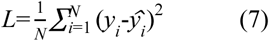

where *N* is the total number of complexes in the training set.

### 4.2. Molecular docking and dynamics simulations of small molecules with pocket proteins

Molecular docking was performed using AutoDock Vina, with protein/ligand preprocessing (water/ion removal, structure optimization, charge assignment) and visualization via PyMOL. Docking regions were defined based on native ligand-binding sites of pocket proteins.

Molecular dynamics (MD) simulations of protein-ligand complexes were conducted with GROMACS. Proteins were parameterized using the CHARMM36 force field, and the TIP3P model was used for solvent. Systems were neutralized with Na⁺/Cl⁻ ions, energy-minimized (nonlinear conjugate gradient method), equilibrated at 300 K/1 bar (1000 ps), and subjected to 100 ns production simulations. Non-bonded interactions were handled with a 10 Å cutoff and PME method, while hydrogen bonds were constrained via the LINCS algorithm (2 fs time step).

Detailed protocols are available in the Supplementary Information.

### 4.3. Affinity Screening and Accurate Determination of Pocket Protein and Small Molecule Interactions

For affinity screening, small molecules were immobilized on the surface of an 8-channel MetaSPR chip via photo-crosslinking. Different pocket proteins were diluted to the same concentration (300 nM), and blank buffer was used for baseline correction. At a flow rate of 40 μL/min, eight different protein solutions were injected simultaneously into the microfluidic system of the chip, and kinetic curves for the association and dissociation phases were monitored under continuous buffer flow.

For accurate affinity determination, the same small molecule was first photo-crosslinked onto the chip. The same protein was then serially diluted from 1152 nM to 72 nM in two-fold steps. Each diluted sample was injected into the microfluidic system of the chip at a flow rate of 40 μL/min, and the association and dissociation kinetics were monitored in real time. All kinetic data were analyzed using WeSPRi 8X software (Xlement, China).

### 4.4. Fabrication of the MetaSPR chip and integration into a POCT platform

The MetaSPR chip was manufactured through a replica-molded nanoimprint process according to our previous report^[37]^. The original mold featuring tapered nanopillar array was patterned on a 12-inch silicon oxide wafer through laser interference lithography and ion etching processes. Following a 12-hour hexylsilane vapor-phase treatment in a vacuum chamber, the surface-engineered model was sequentially coated with the Norland optical adhesive photopolymer and placed with a PET overlay. Following 3-minute UV curing (105 mW/cm²), delamination of the PET substrate featured tapered nanostructures with defined geometric parameters (diameter: 220 nm, height: 500 nm, periodicity: 440 nm). Sequential electron-beam deposition of 5 nm Ti adhesion layer and 60 nm Au plasmonic coating completed the MetaSPR architecture. The final MetaSPR chips were segmented into 5 × 3 mm pieces, allowing for seamless integration with the chip carrier, which was subsequently assembled into the chip cartridge.

The POCT platform incorporated a modular architecture comprising a chip cartridge, pump, optical detection probe, and smartphone app. This app employed Bluetooth (HC-05) for sending commands to the pump and receiving optical data from the photodetector (PD). Pulse-width modulation technology is utilized within these commands to achieve precise regulation of the peristaltic pump’s motor speed, thereby controlling the flow rate through a voltage range of 0-5 V. The PD, integrated into a sophisticated circuit, monitors the intensity variations of 626 nm light on the MetaSPR sensor. These optical signals are relayed to the smartphone app, which processes and visualizes them as dynamic curves to capture molecular interactions.

### 4.5. Ethics approval and consent to participate

The study protocol was approved by the Ethical Review Committee of Jiangsu Provincial Hospital(approval number 2024-008-2).

### 4.6. Statistical analysis

Detailed statistical analysis can be found in the Supplementary Information.

## Author Contributions

Conceptualization: GLL LH XL JG

Methodology: YY HF CJ DL

Investigation: YY CJ HF YL HZ YXZ YYZ YC

Data Curation: YY CJ HF

Formal Analysis: GLL LH HF XL

Supervision: GLL LH XL JG CS

Writing—original draft: YY HF

Writing—review & editing: LH GLL

Funding Acquisition: LH YL GLL CS

Resources: LH

## Acknowledgments

This work was supported by grants from the State Key Laboratory of Microbial Technology Open Projects Fund (2025-19), the Major Program (JD) of Hubei Province (2023BAA011), Fundamental Research Funds for the Central Universities (2021XXJS112), the National Natural Science Foundation of China (82003305), the Fundamental Research Funds for the Central Universities (YCJJ20241402), the Major Project of Guangzhou National Laboratory (GZNL2023A03006), and National Natural Sciences Foundation of China (82202313).

## Data availability statement

The data supporting the findings of this study are available in the supplementary material. The source code and data are available at: https://github.com/popsnake/VD_Pocket.

During the preparation of this work the author(s) used Doubao AI in order to polish the academic English, refine the manuscript expression, and enhance the readability and logic of the article. After using this tool/service, the author(s) reviewed and edited the content as needed and take(s) full responsibility for the content of the published article.

