## Supplemental information for "AI-guided discovery of natural pocket proteins for portable small-molecule sensing"

**This file includes:**

Sections Materials and methods
Figs. S1 to S38
Tables S1 to S6
References

### *1.* Materials and methods

#### *1.1. Materials*

Monoclonal antibodies targeting vitamin B1 (Thiamine) and 25-hydroxyvitamin $D_3$ ($25(OH)D_3$) were obtained from Oukai Biotechnology Co., Ltd. (Nanjing, China). Thiamine, testosterone, and $25(OH)D_3$ were sourced from Beijing Zhongjian Navigation Aid Measurement Technology Co., Ltd. (Beijing, China). Pocket proteins were custom-synthesized by Beijing Sino Biological Inc. (Beijing, China). Thyroid hormones were purchased from Sigma-Aldrich (St. Louis, MO, USA). Hydrogen tetrachloroaurate (III) trihydrate ($HAuCl_4•3H_2O$), trisodium citrate dihydrate ($Na_3C_6H_5O_7•2H_2O$), and sodium dodecyl sulfate were supplied by Beijing Solarbio Science & Technology Co., Ltd. (Beijing, China). SH-PEG-COOH was purchased from Shanghai Aladdin Biochemical Technology Co., Ltd (Shanghai, China). Protein expression for all sequences(1GU3, 1LF9, 4F3K, 1GAI, 2VZR, 1NLI,1GNY, 1) was provided by Sangon Biotech (Shanghai, China). All reagents were used as received without further purification.

**Table S1. Sequences of Thiamine Pocket Proteins.**

| No. | PDB | Sequence |
|---|---|---|
| 1 | **1J5I** | AAPTATVTPSSGLSDGTVVKVAGAGLQAGTAYDVGQCAWVDTGVLACNP<br>ADFSSVTADANGSASTSLTVRRSFEGFLFDGTRWGTVDCTTAACQVGLSDA<br>AGNGPEGVAISFN |
| 2 | **1HEE** | ARSTNTFNYATYHTLDEIYDFMDLLVAQHPELVSKLQIGRSYEGRPIYVLKFST<br>GGSNRPAIWIDLGIHSREWITQATGVWFAKKFTENYGQNPSFTAILDSMDIFL<br>EIVTNPNGFAFTHSENRLWRKTRSVTSSSLCVGVDANRNWDAGFGKAGASS<br>SPCSETYHGKYANSEVEVKSIVDFVKNHGNFKAFLSIHSYSQLLLYPYGYTTQS<br>IPDKTELNQVAKSAVAALKSLYGTSYKYGSIITTIYQASGGSIDWSYNQGIKYSF<br>TFELRDTGRYGFLLPASQIIPTAQETWLGVLTIMEHTVNN |
| 3 | **1JJK** | VTNSPVVVALDYHNRDDALAFVDKIDPRDCRLKVGKEMFTLFGPQFVRELQ<br>QRGFDIFLDLKFHDIPNTAAHAVAAAADLGVWMVNVHASGGARMMTAAR<br>EALVPFGKDAPLLIAVTVLTSMEASDLVDLGMTLSPADYAERLAALTQKCGLD<br>GVVCSAQEAVRFKQVFGQEFKLVTPGIRPQGSEAGDQRRIMTPEQALSAGV<br>DYMVIGRPVTQSVDPAQTLKAINASLQ |
| 4 | **4F3K** | MKIGIIGAMEEEVTLLRDKIDNRQTITLGGCEIYTGQLNGTEVALLKSGIGKVAA<br>ALGATLLLEHCKPDVIINTGSAGGLASTLKVGDIVVSDETRYHDADVTAFGYE<br>YGQLPGCPAGFKADDKLIAAAESCIRELNLNAVRGLIVSGDAFINGSVGLAKIR<br>HNFPDAVAVEMEATAIAHVCHNFNVPFVVVRAISDVADQQSHLSFDEFLAV<br>AAKQSTLMVETLVQKLA |
| 5 | **4KLB** | APAAVDWRARGAVTAVKDQGQCGSCWAFSAIGNVECQWFLAGHPLTNLS<br>EQMLVSCDKTDSGCSGGLMNNAFEWIVQENNGAVYTEDSYPYASGEGISPP<br>CTTSGHTVGATITGHVELPQDEAQIAAWLAVNGPVAVAVDASSWMTYTGG<br>VMTSCVSEQLDHGVLLVGYNDSAAVPYWIIKNSWTTQWGEEGYIRIAKGSN<br>QCLVKEEASSAVVG |
| 6 | **1GNY** | GNVVIEVDMANGWRGNASGSTSHSGITYSADGVTFAALGDGVGAVFDIAR<br>PTTLEDAVIAMVVNVSAEFKASEANLQIFAQLKEDWSKGEWDCLAGSSELTA<br>DTDLTLTCTIDEDDDKFNQTARDVQVGIQAKGTPAGTITIKSVTITLAQEA |
| 7 | **4Q2K** | CGVPAIQPVLIVNGEEAVPGSWPWQVSLQDKTGFHFCGGSLINENWVVTA<br>AHCGVTTSDVVVAGEFDQGSSSEKIQKLKIAKVFKNSKNDITLLKLSTAASFSQ<br>TVSAVCLPSASDDFAAGTTCVTTGWGLTRYANTPDRLQQASLPLLSNTNCK<br>KYWGTKIKDAMICAGASGVSSCMGDSGGPLVCKKNGAWTLVGIVSWGSST<br>CSTSTPGVYARVTALVNWVQQTLAAN |
| 8 | **1GAI** | ATLDSWLSNEATVARTAILNNIGADGAWVSGADSGIVVASPSTDNPDYFYT<br>WTRDSGLVIKTLVDLFRNGDTDLLSTIEHYISSQAIIQGVSNPSGDLSSGGLGE<br>PKFNVDETAYTGSWGRPQRDGPALRATAMIGFGQWLLDNGYTSAATEIVW<br>PLVRNDLSYVAQYWNQTGYDLWEEVNGSSFFTIAVQHRALVEGSAFATAVG<br>SSCSWCDSQAPQILCYLQSFWTGSYILANFDSSRSGKDTNTLLGSIHTFDPEA<br>GCDDSTFQPCSPRALANHKEVVDSFRSIYTLNDGLSDSEAVAVGRYPEDSYY<br>NGNPWFLCTLAAAEQLYDALYQWDKQGSLEITDVSLDFFKALYSGAATGTY<br>SSSSSTYSSIVSAVKTFADGFVSIVETHAASNGSLSEQFDKSDGDELSARDLTW |

| | |
|---|---|
| | SYAALLTANNRRNSVVPPSWGETSASSVPGTCAATSASGTYSSVTVTSWPSIVATG |

**Table S2. Sequences of 25(OH)$D_3$ Pocket Proteins.**

| No. | PDB | Pocket Name |
|---|---|---|
| 1 | **4FE9** | GQLTATFTAGKDASIVMDAATADSVEIAKLSSTTAEEGSKIAVNSLTLNENHTI<br>PFSMTEDHVFKVALAQLDSVTQEAYKSRASVVRELKISINASAVTPSGEGIQL<br>VGNEVSITLQPATTPAVDPDGYYIVGDFTGWDGNSAQQMKKDALDENLYIL<br>EAEIESTSNFKIFPASAINGNDIDWTKALGSSVDGDDSGDNFVSWTNAGAIN<br>TALDGKIKISFDAFNYRFTVKDNSAPTELYMTGSAYNWGTPAGDPNAWKAL<br>VPVNGTKGTFWGIFYFAANDQVKFAPQANWGNDFGFVDAISQESKDLAGL<br>SDEGGNLKVGIAGWYLVYVSVIGDDKVIEFEKPNVYLMGDTSYNGWDAQLV<br>EQDLFTVPGTADGEFVSPAFLKDGAVRICVNPKAVSAGDWWKTEFIIFDGQI<br>AYRGNGGDQAAVQGKTGQKVYLNFGNGTGRIE |
| 2 | **2JDL** | MVSFRLSGATSSSYGVFISNLRKALPNERKLYDIPLLRSSLPGSQRYALIHLTNY<br>ADETISVAIDVTNVYIMGYRAGDTSYFFNEASATEAAKYVFKDAMRKVTLPYS<br>GNYERLQTAAGKIRENIPLGLPALDSAITTLFYYNANSAASALMVLIQSTSEAA<br>RYKFIEQQIGKRVDKTFLPSLAIISLENSWSALSKQIQIASTNNGQFESPVVLIN<br>AQNQRVTITNVDAGVVTSNIALLLNRNNMA |
| 3 | **1JJK** | VTNSPVVVALDYHNRDDALAFVDKIDPRDCRLKVGKEMFTLFGPQFVRELQ<br>QRGFDIFLDLKFHDIPNTAAHAVAAAADLGVWMVNVHASGGARMMTAAR<br>EALVPFGKDAPLLIAVTVLTSMEASDLVDLGMTLSPADYAERLAALTQKCGL<br>DGVVCSAQEAVRFKQVFGQEFKLVTPGIRPQGSEAGDQRRIMTPEQALSAG<br>VDYMVIGRPVTQSVDPAQTLKAINASLQ |
| 4 | **1LF9** | SIKIDRFNNISAVNGPGEEDTWASAQKQGVGTANNYVSKVWFTLANGAISE<br>VYYPTIDTADVKEIKFIVTDGKSFVPDETKDAISKVEKFTDKSLGYKLVNTDKKG<br>RYRITKDIFTDVKRNSLIMKAKFEALEGSIHDYKLYLAYDPHIKNQGSYNEGYVI<br>KANNNEMLMAKRDNVYTALSSNIGWKGYSIGYYKVNDIMTDLDENKQMT<br>KHYDSARGNIIEGAEIDLTKNSEFEIVLSFGQSDSEAAKTALETLGEDYNNLKN<br>NYIDEWTKYCNTLNNFNGKANSLYYNSMMILKASEDKTNKGAYIASLSIPW<br>GDGQRDDNTGGYHLVWSRDLYHVANAFIAAGDVDSANRSLDYLAKVVKD<br>NGMIPQNTWISGKPYWTGIQLDEQADPIILSYRLKRYDLYDSLVKPLADFIIKI<br>GPKTGQERWEEIGGYSPATMAAEVAGLTCAAYIAEQNKDYESAQKYQEKAD<br>NWQKLIDNLTYTENGPLGNGQYYIRIAGLSDPDADFMINIANGGGVYDQKE<br>IVDPSFLELVRLGVKSADDPKILNTLKVVDSTIKVDTPKGPSWYRYNHDGYGE<br>PSKTELYHGAGKGRLWPLLTGERGMYEIAAGKDATPYVKAMEKFANEGGIIS<br>EQVWEDTGLPTDSASPLNWAHAEYVILFASNIEHKVLDMPDIVYKRYVA |
| 5 | **4CSD** | MSSVQTAATSWGTVPSIRVYTANNGKITERCWDGKGWYTGAFNEPGDNV<br>SVTSWLVGSAIHIRVYASTGTTTTEWCWDGNGWTKGAYTS-----<br>DQTAATSWGTVPSIRVYTANNGKITERCWDGKGWYTGAFNEPGDNVSVTS<br>WLVGSAIHIRVYASTG-<br>TTTEWCWDGNGWTKGAYTSSTVPGDQTAATSWGTVPSIRVYTANNGKITE<br>RCWDGKGWYTGAFNEPGDNVSVTSWLVGSAIHIRVYASTGTTTTEWCWD<br>GNGWTKGAYTAT |

| | | |
|---|---|---|
| 6 | **1ADD** | TPAFNKPKVELHVHLDGAIKPETILYFGKKRGIALPADTVEELRNIIGMDKPLSL<br>PGFLAKFDYYMPVIAGCREAIKRIAYEFVEMKAKEGVVYVEVRYSPHLLANSK<br>VDPMPWNQTEGDVTPDDVVDLVNQGLQEGEQAFGIKVRSILCCMRHQPS<br>WSLEVLELCKKYNQKTVVAMDLAGDETIEGSSLFPGHVEAYEGAVKNGIHRT<br>VHAGEVGSPEVVREAVDILKTERVGHGYHTIEDEALYNRLLKENMHFEVCPW<br>SSYLTGAWDPKTTHAVVRFKNDKANYSLNTDDPLIFKSTLDTDYQMTKKDM<br>GFTEEEFKRLNINAAKSSFLPEEEKKELLERLYREYQ |
| 7 | **1JYS** | MKIGIIGAMEEEVTLLRDKIENRQTISLGGCEIYTGQLNGTEVALLKSGIGKVAA<br>ALGATLLLEHCKPDVIINTGSAGGLAPTLKVGDIVVSDEARYHDADVTAFGYE<br>YGQLPGCPAGFKADDKLIAAAEACIAELNLNAVRGLIVSGDAFINGSVGLAKI<br>RHNFPQAIAVEMEATAIAHVCHNFNVPFVVVRAISDVADQ----<br>SFDEFLAVAAKQSSLMVESLVQKLA |
| 8 | **1GAI** | ATLDSWLSNEATVARTAILNNIGADGAWVSGADSGIVVASPSTDNPDYFYT<br>WTRDSGLVIKTLVDLFRNGDTDLLSTIEHYISSQAIIQGVSNPSGDLSSGGLGE<br>PKFNVDETAYTGSWGRPQRDGPALRATAMIGFGQWLLDNGYTSAATEIVW<br>PLVRNDLSYVAQYWNQTGYDLWEEVNGSSFFTIAVQHRALVEGSAFATAV<br>GSSCSWCDSQAPQILCYLQSFWTGSYILANFDSSRSGKDTNTLLGSIHTFDPE<br>AGCDDSTFQPCSPRALANHKEVVDSFRSIYTLNDGLSDSEAVAVGRYPEDSY<br>YNGNPWFLCTLAAAEQLYDALYQWDKQGSLEITDVSLDFFKALYSGAATGT<br>YSSSSSTYSSIVSAVKTFADGFVSIVETHAASNGSLSEQFDKSDGDELSARDLT<br>WSYAALLTANNRRNSVVPPSWGETSASSVPGTCAATSASGTYSSVTVTSWP<br>SIVATG |

### *1.2. High-Efficiency Screening of Thiamine- and 25(OH)$D_3$-Binding Pocket Proteins using the PLCA model*

The screening protocol was as follows: protein structures were obtained from the PDBbind 2016 database and preprocessed using PyMOL to remove crystallographic water molecules, ions, and extraneous ligands, thereby isolating the putative binding pockets. The 3D structures of the small molecules, Thiamine (PubChem CID: 1130) and 25(OH)$D_3$ (PubChem CID: 5283731), were retrieved from the PubChem database. The **PLCA** model utilizes a dual-graph encoder architecture to represent both the protein pocket and the small molecule as graphs. Its core component, the EGMN, performs joint transformations on both atomic features and 3D coordinates through equivariant operations. This enables the model to explicitly account for pocket flexibility, such as side-chain rotations and backbone adjustments, which are critical for binding. The model was pre-trained using two self-supervised tasks—an atom-level prompt-based denoising generative task and a conformation-level snapshot ordering task—endowing it with a robust understanding of binding-related dynamics. we repurposed **PLCA** for pocket protein screening, where the target small molecules (Thiamine and 25(OH)$D_3$) were fixed as probes to scan the entire protein database for potential high-affinity binders. The binding score for each protein-small molecule pair was calculated, leading to the initial selection of 40 candidate pocket proteins for Thiamine and 40 for 25(OH)$D_3$. This candidate set was subsequently refined by removing redundant sequences through clustering analysis based on sequence similarity, yielding a final, non-redundant list of pocket proteins for experimental validation.

### *1.3. Molecular docking and dynamics simulations of small molecules with pocket proteins*

We employed the AutoDock Vina software for molecular docking experiments between proteins and small molecules. AutoDock Vina is a widely used molecular docking tool that efficiently predicts the binding modes and affinities of small molecule ligands with macromolecular targets. Additionally, PyMOL software was utilized for the visualization and preprocessing of molecular structures. Using PyMOL, we performed several preprocessing steps on the remaining structures, including the removal of water molecules, ions, and ligands, as well as the elimination of redundant structures in multimers. The PDB files were then converted into PDBQT files recognizable by Vina, with hydrogen atoms added simultaneously. The small molecules (25(OH)$D_3$ or Thiamine) were optimized to ensure it adopted a low-energy conformation. Hydrogen atoms were also added to the ligand, and appropriate charges were assigned to enhance the accuracy of the docking process. The active sites of the proteins were located based on the binding conditions of the original ligands in the pocket proteins, and the docking region parameters were appropriately configured. Subsequent docking calculations were performed with reference to this configuration.

Molecular dynamics simulations of the protein-small molecule complexes were conducted using the GROMACS software package. The protein structures obtained from the AutoDock Vina were preprocessed using PyMOL, which included the removal of crystallographic waters and the addition of missing hydrogen atoms, ultimately generating the protein structure files for simulation. Subsequently, the pdb2gmx tool from GROMACS was utilized to parameterize the protein with the CHARMM36 force field, and the TIP3P model was selected as the water force field for the solvent. To ensure system electroneutrality, appropriate amounts of $Na^+$ and $Cl^-$ ions were added to the system using the genion tool.

In terms of system construction, the editconf tool was used to define a cubic box centered on the structure, which was then filled with solvent water molecules using the solvate tool. During the energy minimization phase, the system was optimized using the nonlinear conjugate gradient method, with a force tolerance set at 1000 kJ/mol·nm. Subsequently, the system was equilibrated under constant temperature (300 K) and pressure (1 bar) conditions for approximately 1000 ps, with temperature and pressure coupling performed using the Berendsen algorithm. For non-bonded interactions, the cutoff distance between atom pairs was set at 10 Å, and long-range electrostatic interactions were handled using the Particle Mesh Ewald (PME) method. Additionally, the LINCS algorithm was employed to constrain hydrogen bonds, thereby enhancing simulation efficiency. The time step for all simulations was set at 2 fs.

After completing the equilibration simulations, the production phase of the MD simulations was carried out for a duration of 100 ns. During the simulations, trajectory and energy files were regularly outputted for subsequent analysis. Through these steps, we successfully constructed and simulated the protein-small molecule ligand complex systems, providing a foundation for in-depth studies of their dynamic behavior and interaction mechanisms.

*1.4. Small molecules modification using photo-crosslinking technology*

The MetaSPR chip was sequentially rinsed with ultrapure water and anhydrous ethanol. After ensuring complete evaporation of residual liquids, 2 μL of 1 mM SH-PEG-COOH solution was dispensed into each well and incubated overnight at 4°C. Following incubation, the liquid was discarded, and the wells were washed three times with ultrapure water before drying in a 37°C oven for 8 min. 20 μL EDC/NHS (3-ethylcarbodiimide hydrochloride/ N-hydroxysuccinimide) mixture (400 mM/100 mM, 1:1, v/v) was added to the chip and incubated at 37°C for 20 min. After discarding the reaction solution, the wells were washed twice with ultrapure water and dried in a 37°C oven. Then, 1 μL of 10 mM photocrosslinking agent was added to the chip and incubated at room temperature for 4 h in the dark condition. Subsequently, the wells were washed twice with water and dried at 37°C. 1.7 μL of 10 mM small molecule solution（Thiamine or 25(OH)$D_3$） was dispensed into chip under dark conditions. The modified chip was then exposed to light in a photocrosslinking instrument for 15 min. Then, the wells were washed once with ethanol and three times with ultrapure water. 20 μL of blocking buffer containing 0.05% fish gelatin and 0.5% casein was added on the chip at 25°C for 1h. The blocking solution was discarded, and the modified chip was dried at 25°C followed by storage at 4°C until use.

*1.5. AuNP synthesis and labeling with pocket proteins*

Colloidal AuNPs were synthesized via chemical reduction of $HAuCl4\cdot 3H_2O$ by $Na_3C_6H_5O_7\cdot 2H_2O$, adapting a reported method[1]. Briefly, boiling HAuCl4 (50 mL, 0.02 wt%) solution was rapidly mixed with $Na_3C_6H_5O_7$ solution (0.8 mL, 1 wt%) under continuous stirring. The solution color transitioned from yellow through black to red within minutes, indicating AuNPs formation. These synthesized AuNPs were subsequently functionalized with pocket protein for competitive detection. For labeling, the pH of colloidal AuNPs (1.5 mL) was adjusted to 8.0 using 1 M Tris (pH 9.27). Pocket protein was then introduced at final concentrations of 917 ppb, followed by 15 min incubation. Subsequent blocking employed by BSA (final concentration 1 wt%) with a further 15 min incubation. Finally, the functioned AuNPs were isolated by centrifugation (5400 × g, 20 min), vacuum-dried, and stored at 4°C until required.

*1.6. Multiplexed small molecules detection using pocket protein*

Prior to small molecule detection, the modified chip was integrated into chip card and the AuNPs was redissolved in 500 μL of Tris buffer (20 mM Tris-HCl, pH 8.0, supplemented with 0.3 wt% sucrose and 0.05 wt% PEG 20000). We prepared the real samples for detection by first centrifuging blood to collect serum and then diluting it with PBST. These samples were subsequently used for the multiplexed quantitative detection of small molecules. For curve fitting or analyte detection, 100 μL of the corresponding solution (Thiamine standards: 0–2560 ppt; 25(OH)$D_3$ standards: 0–3200 ppt, both serially diluted two-fold; or analyte samples) was mixed with 6 μL of 0.06 wt% AuNPs and injected onto the chip at 25 μL/min, with sequential injection starting from the highest concentration. After each detection cycle, the sensor was regenerated by flowing 200 μL of 0.5 wt% sodium dodecyl sulfate through the system at 80 μL/min to remove bound analytes.

*1.7. Statistical analysis*

Statistical analyses were performed using SPSS 19.0 (SPSS, USA) and MedCalc 18.11.6 (MedCalc Software, Belgium). All samples were tested in triplicate, and the data were normalized and presented as mean ± standard deviation. The accuracy of the pocket protein sensor and the commercial ELISA method was compared using receiver operating characteristic curve (ROC) analysis on real samples. Additionally, the Passing-Bablok regression method was employed to assess the correlation coefficient. A p-value of less than 0.05 was considered statistically significant.

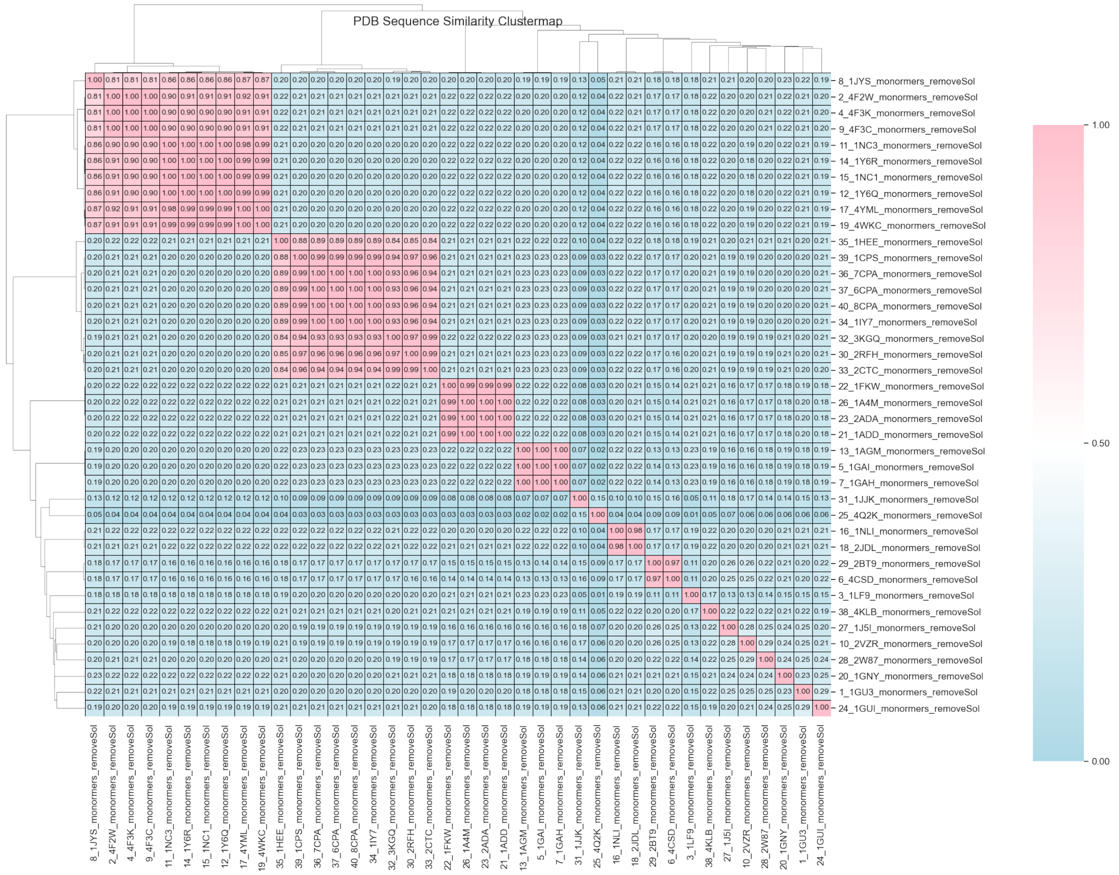


**Fig. S1**. Similarity and clustering analysis results of the candidate protein sequences for Thiamine.

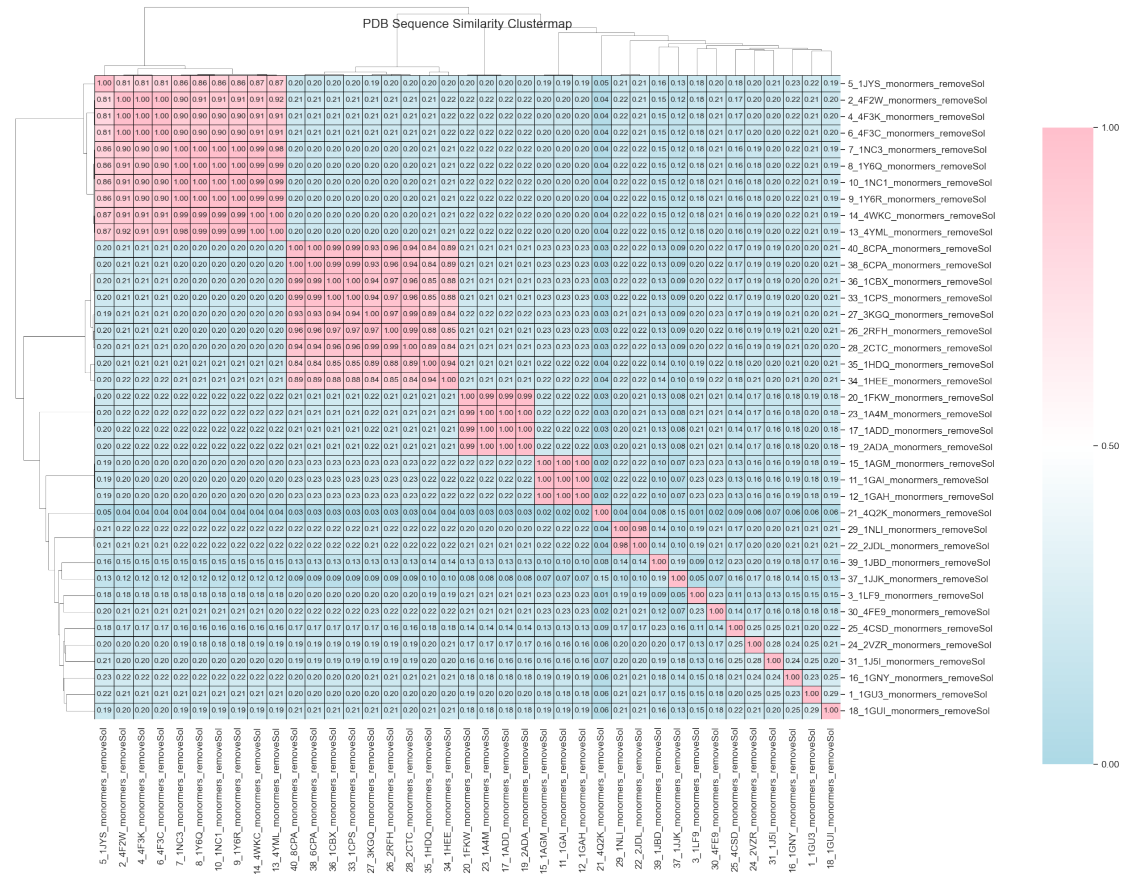

**Fig.S2**. Similarity and clustering analysis results of the candidate protein sequences for 25(OH)$D_3$.

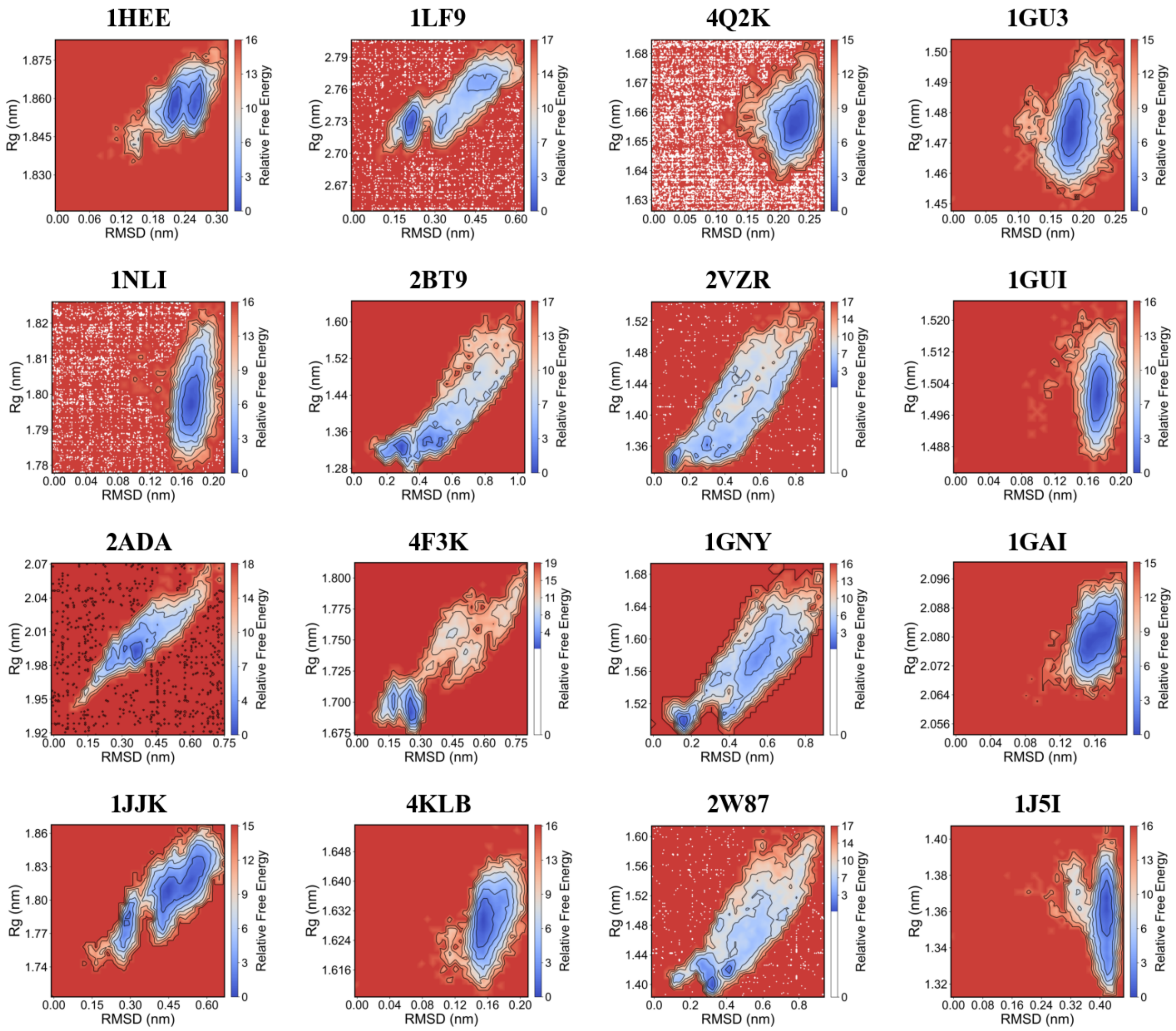


**Fig. S3.** 2D projections of the free energy landscapes of Thiamine.

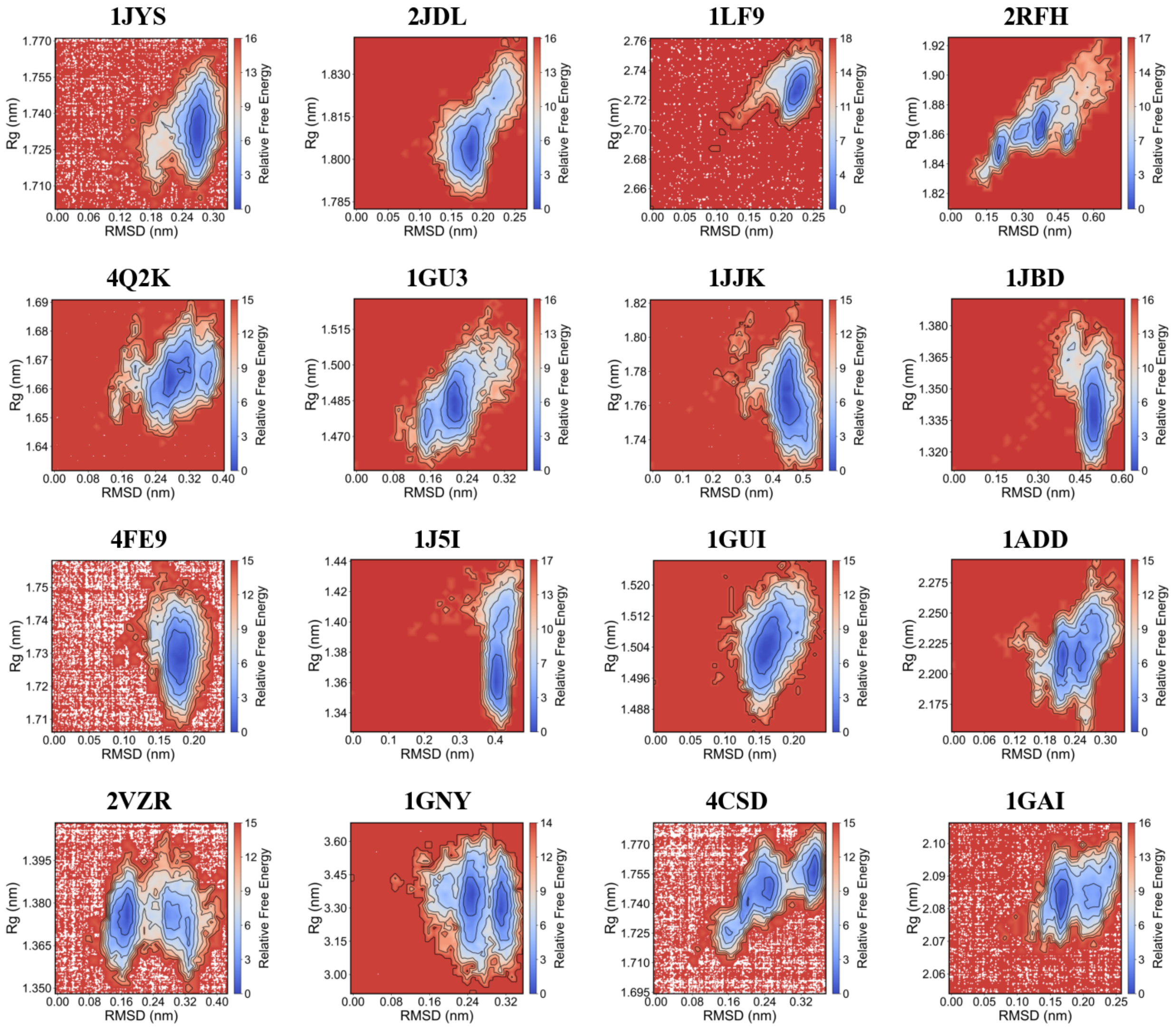


**Fig. S4.** 2D projections of the free energy landscapes of Thiamine.

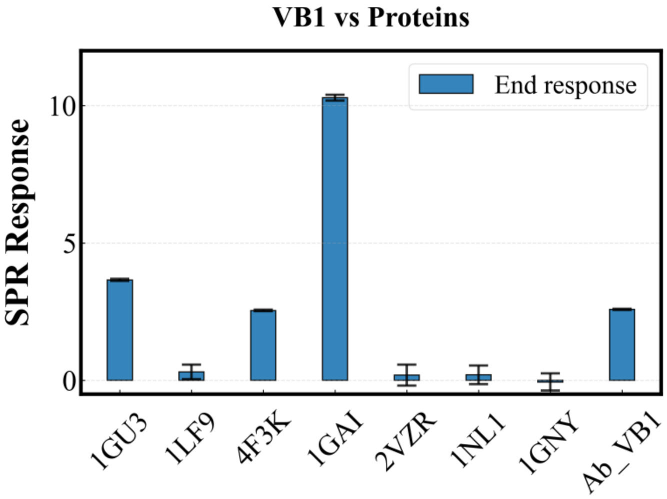


**Fig. S5.** The MetaSPR endpoint signal response comparison of Thiamine with 8 candidate proteins (all at 300 nM protein concentration)

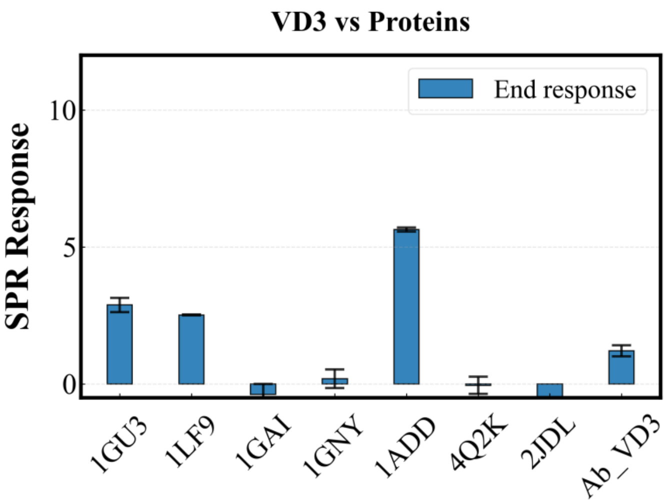


**Fig. S6.** The MetaSPR endpoint signal response comparison of 25(OH)$D_3$ with 8 candidate proteins (all at 300 nM protein concentration)

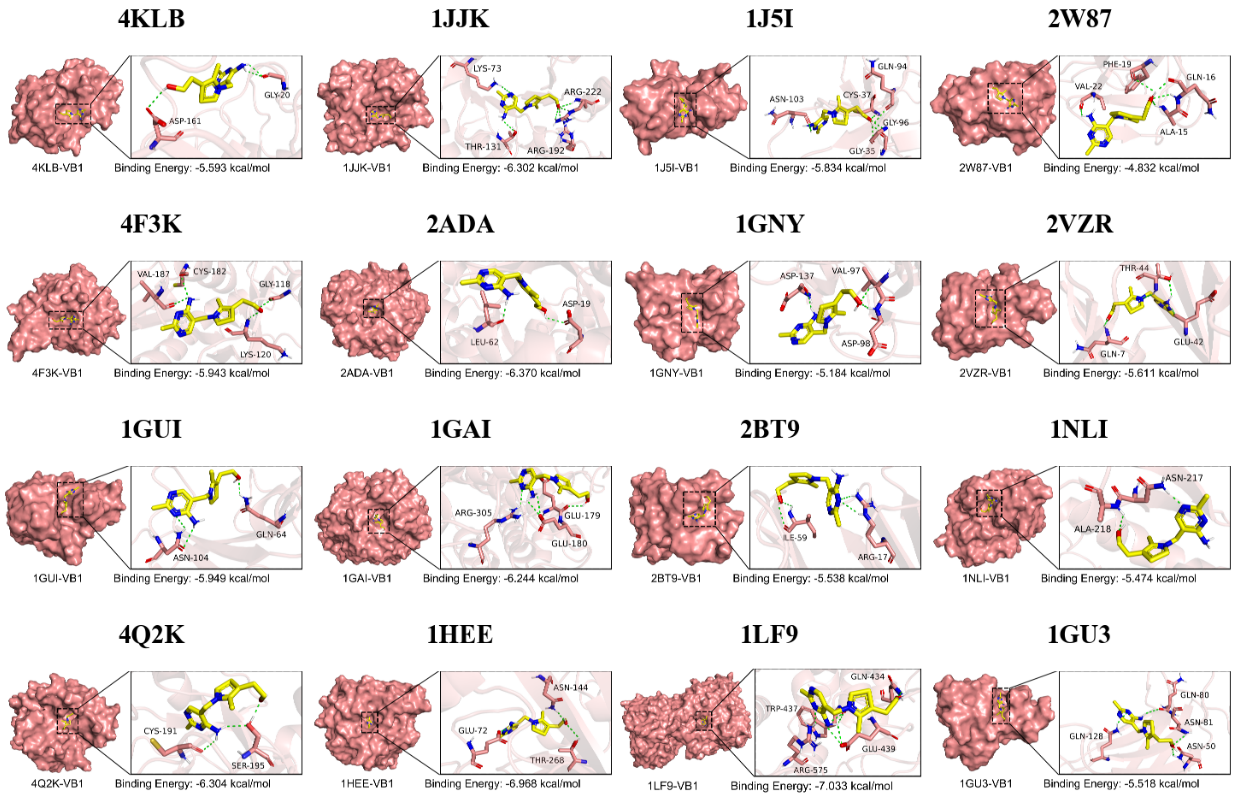


**Fig. S7.** Docking results for Thiamine.

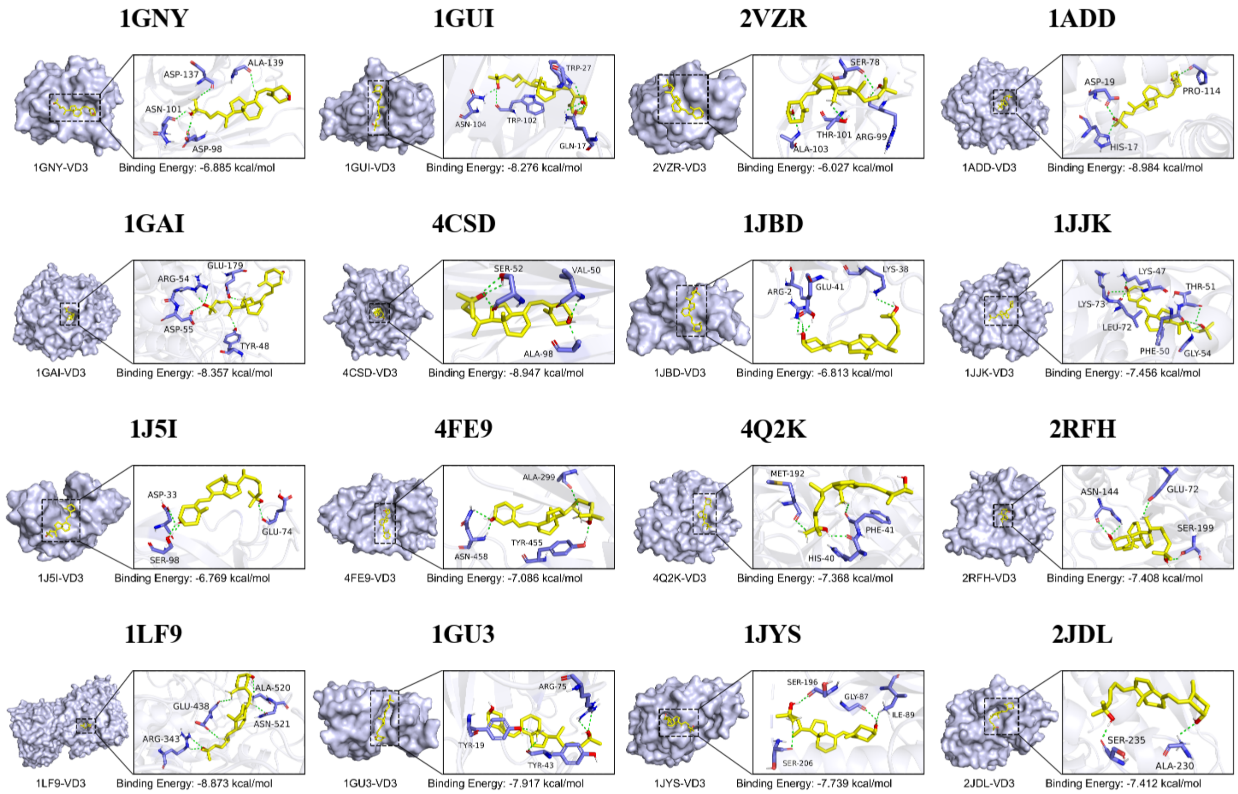


**Fig. S8.** Docking results for 25(OH)$D_3$.

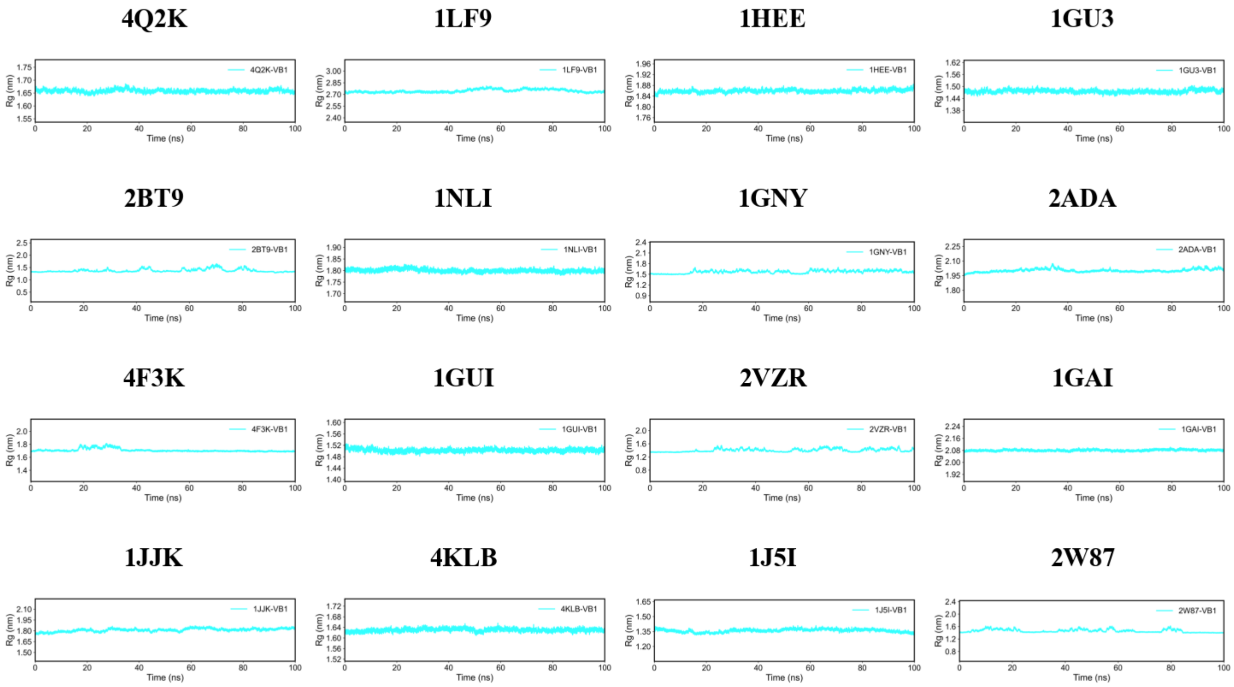


**Fig. S9.** The Radius of Gyration (Rg) of Thiamine with the binding pocket protein.

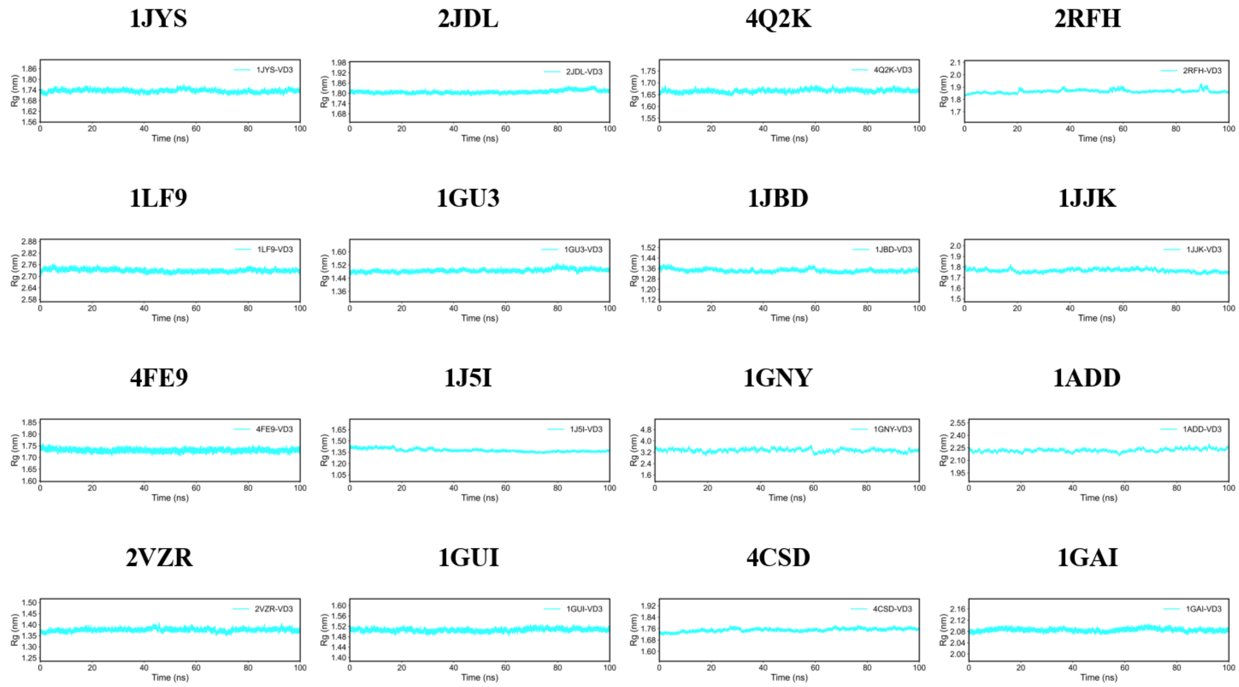


**Fig. S10.** The Radius of Gyration (Rg) of 25(OH)$D_3$ with the binding pocket protein.

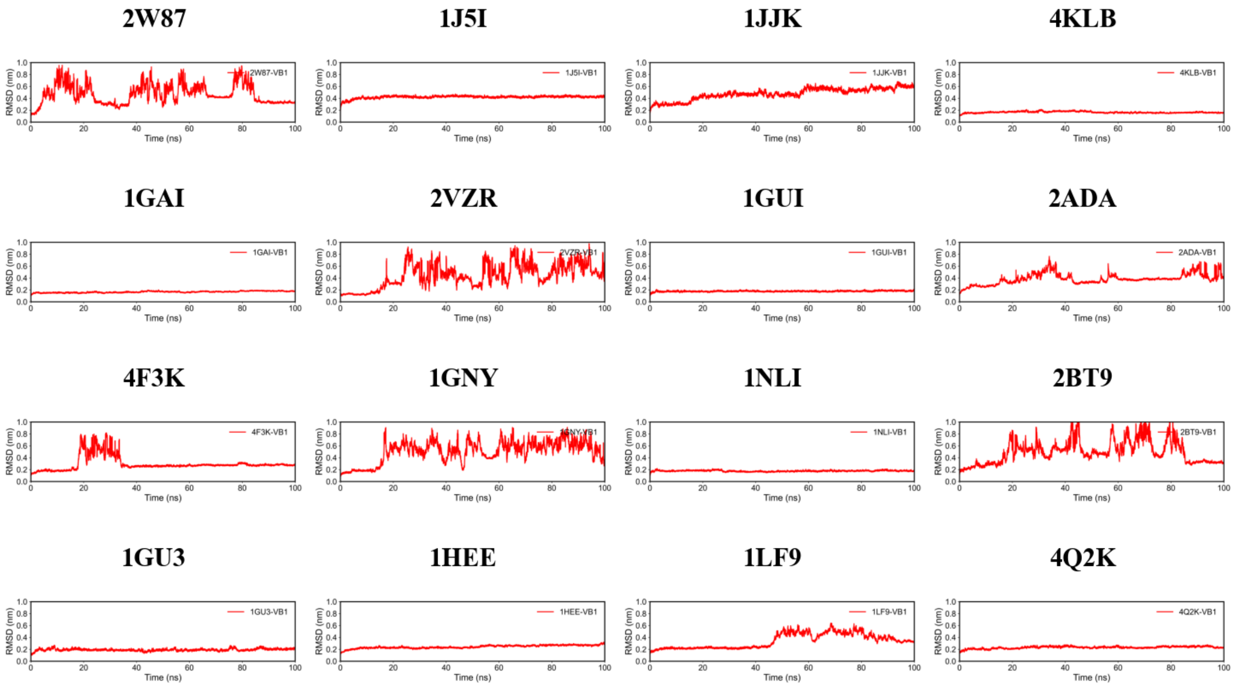


**Fig. S11.** The Root Mean Square Deviation (RMSD) curve of V B in complex with the binding pocket protein.

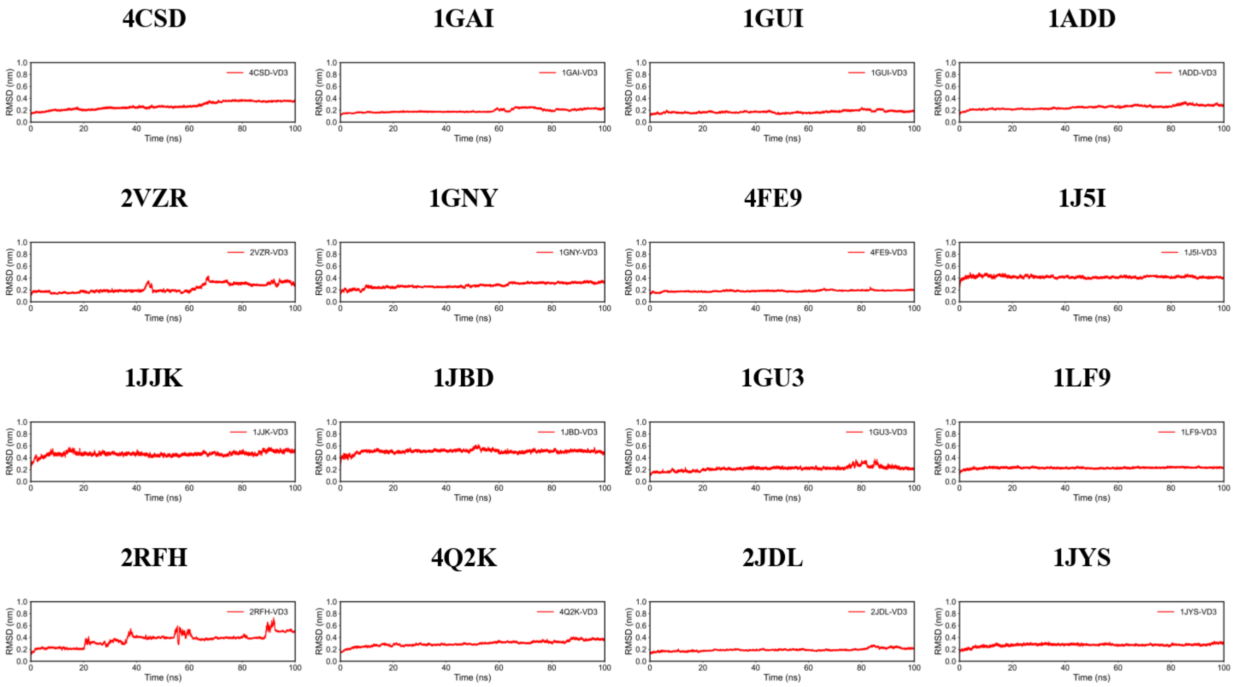


**Fig. S12.** The Root Mean Square Deviation (RMSD) curve of 25(OH)$D_3$ in complex with the binding pocket protein.

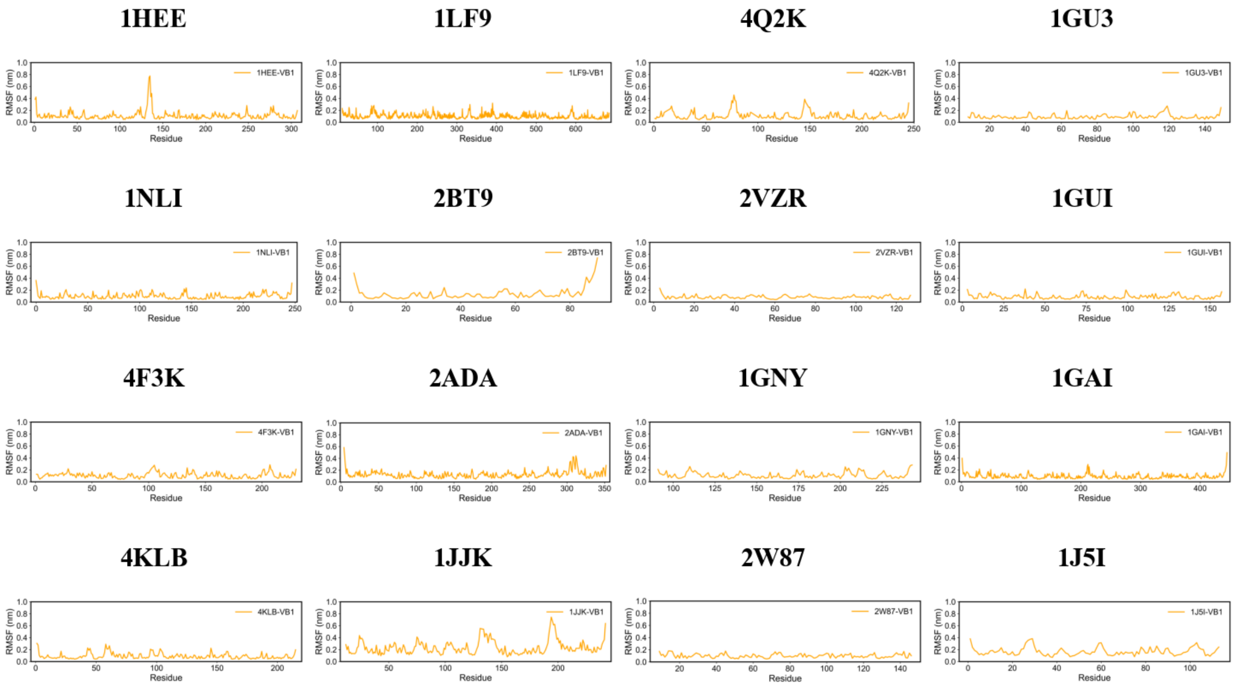


**Fig. S13.** The Root Mean Square Fluctuation (RMSF) of Thiamine in complex with the binding pocket protein.

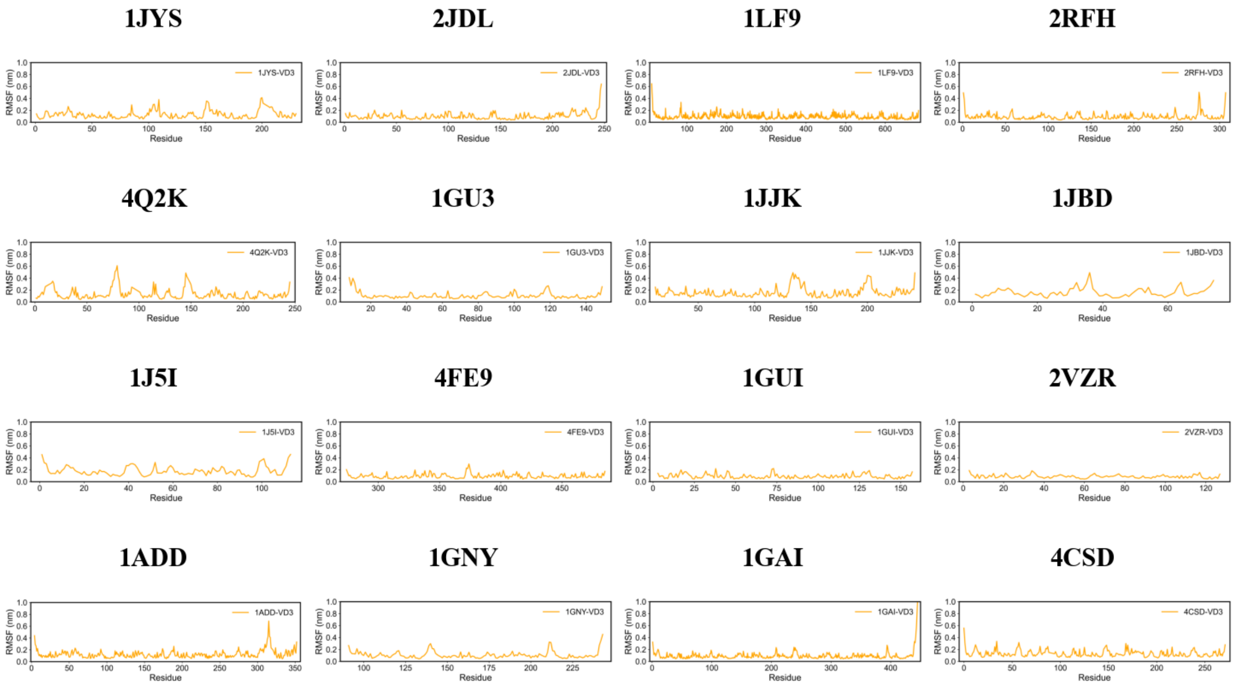


**Fig. S14.** The Root Mean Square Fluctuation (RMSF) of 25(OH)$D_3$ in complex with the binding pocket protein.

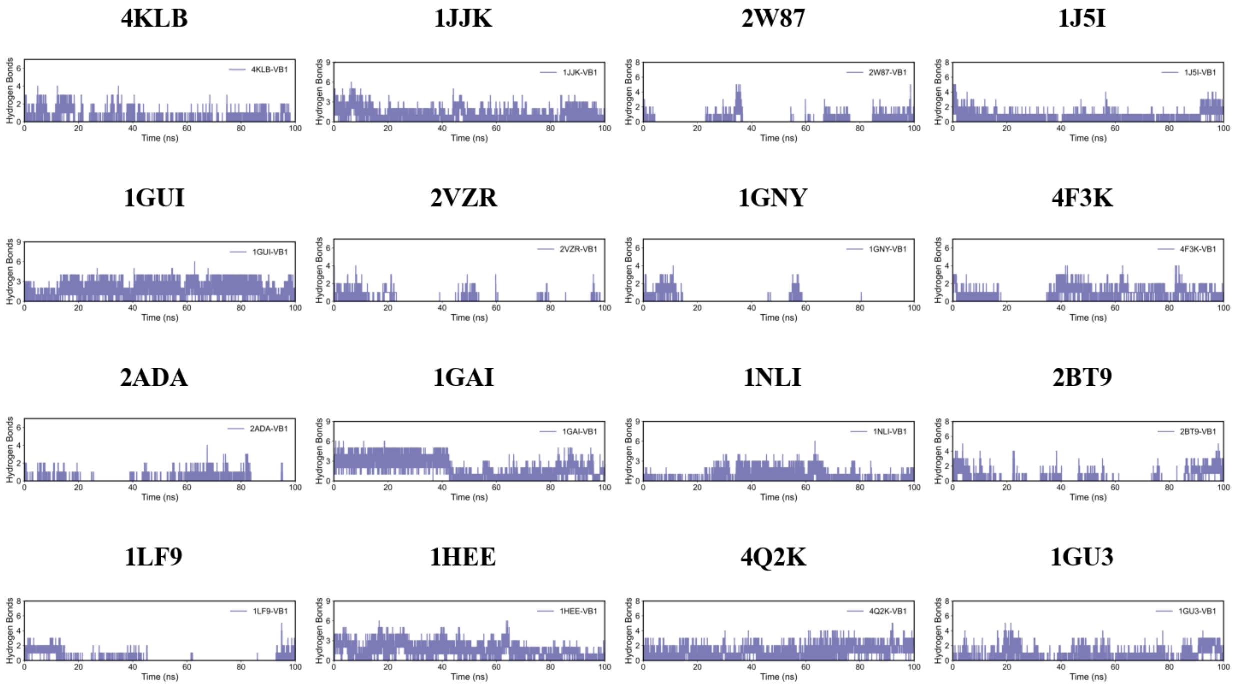


**Fig. S15.** The Hydrogen bonds of VB in complex with the binding pocket protein.

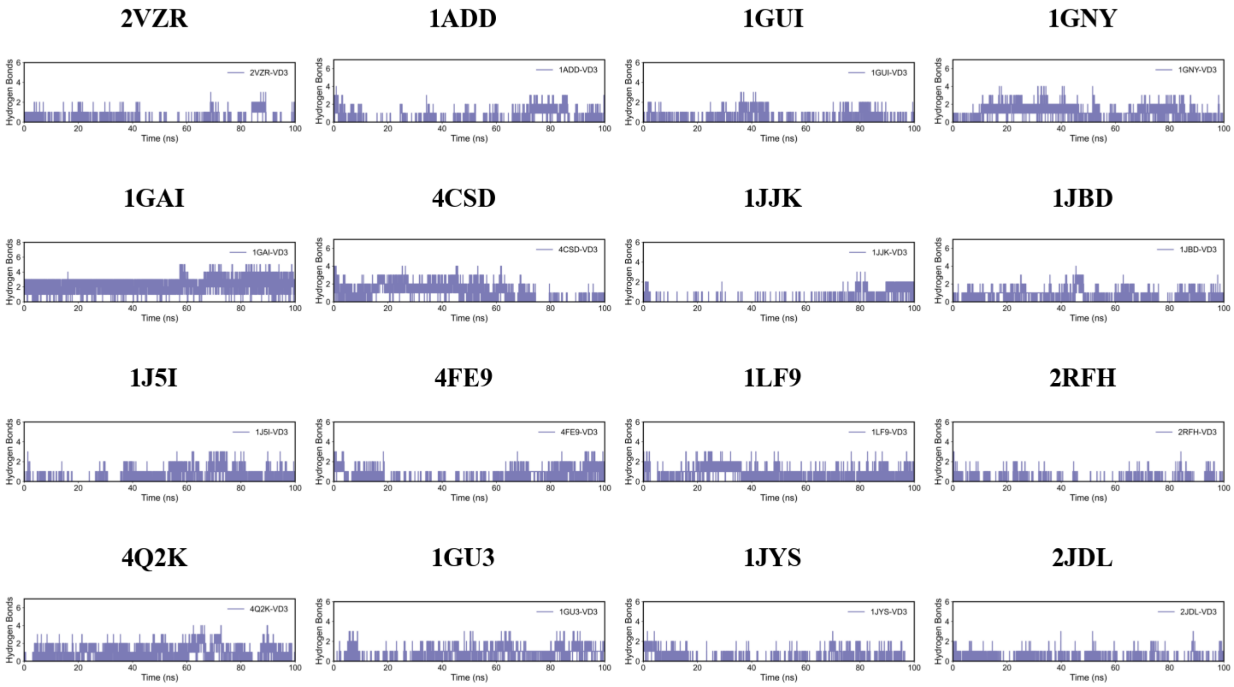


**Fig. S16.** The Hydrogen bonds of VB in complex with the binding pocket protein.

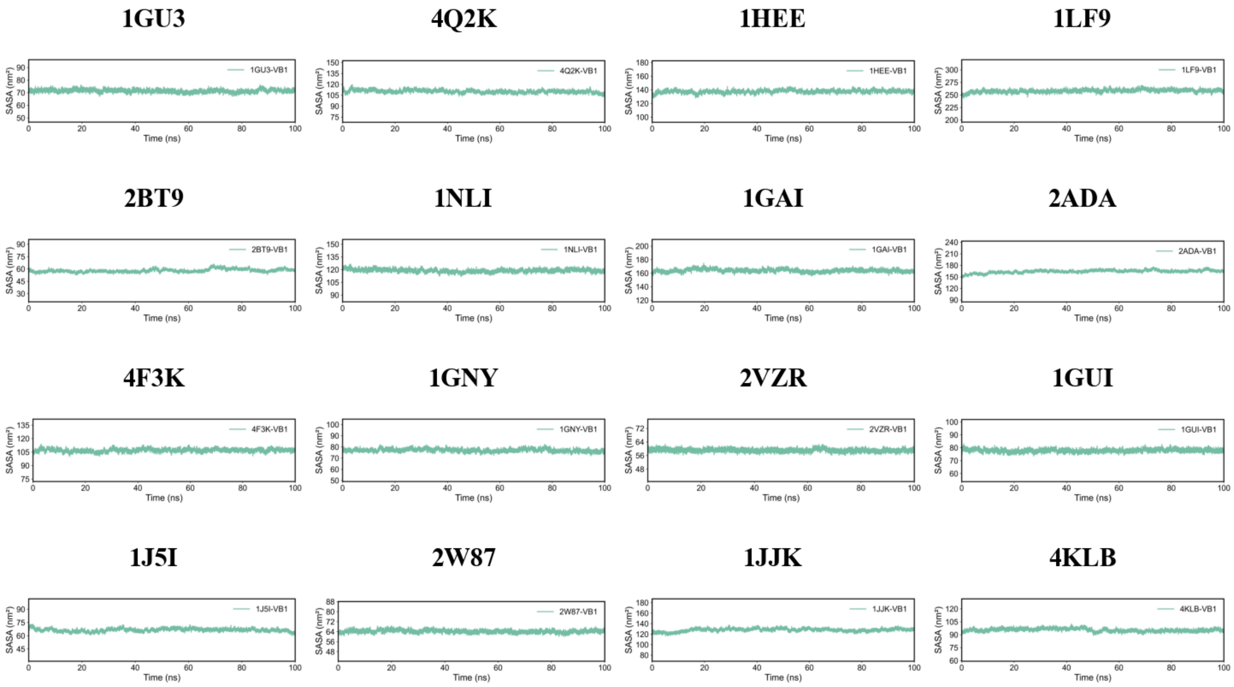


**Fig. S17.** The Solvent Accessible Surface Area (SASA) of Thiamine in complex with the binding pocket protein.

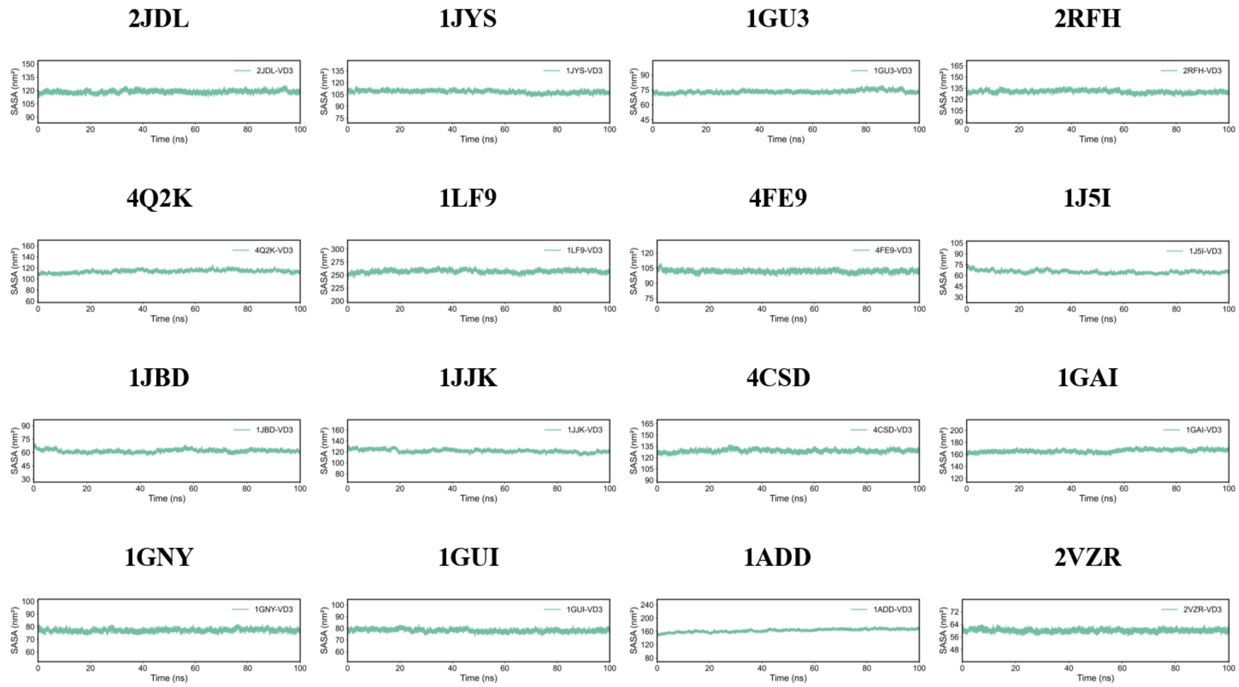


**Fig. S18.** The Solvent Accessible Surface Area (SASA) of 25(OH)$D_3$ in complex with the binding pocket protein.

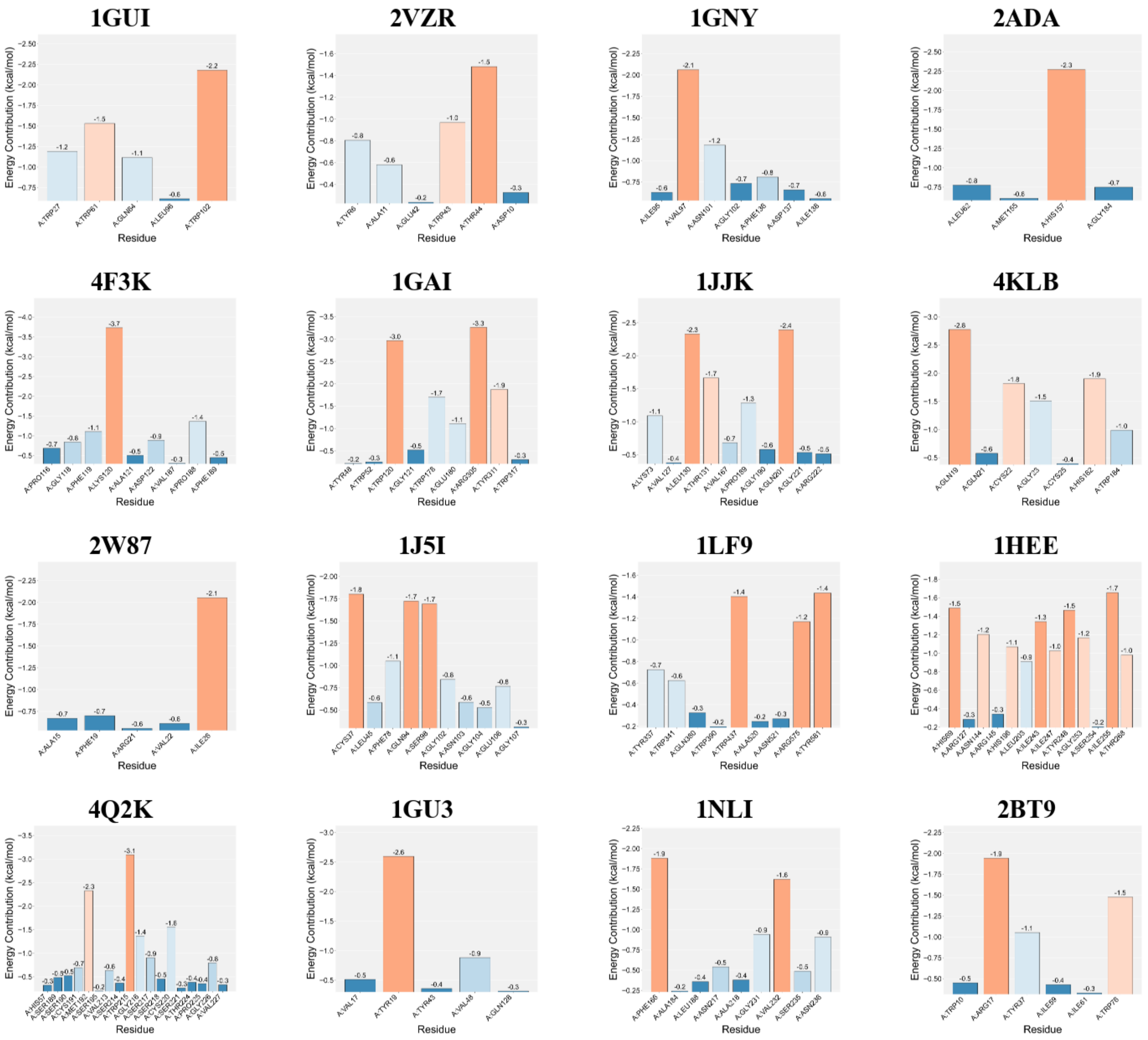


**Fig. S19.** The energy contribution of amino acid residues in the Thiamine complex with the binding pocket protein.

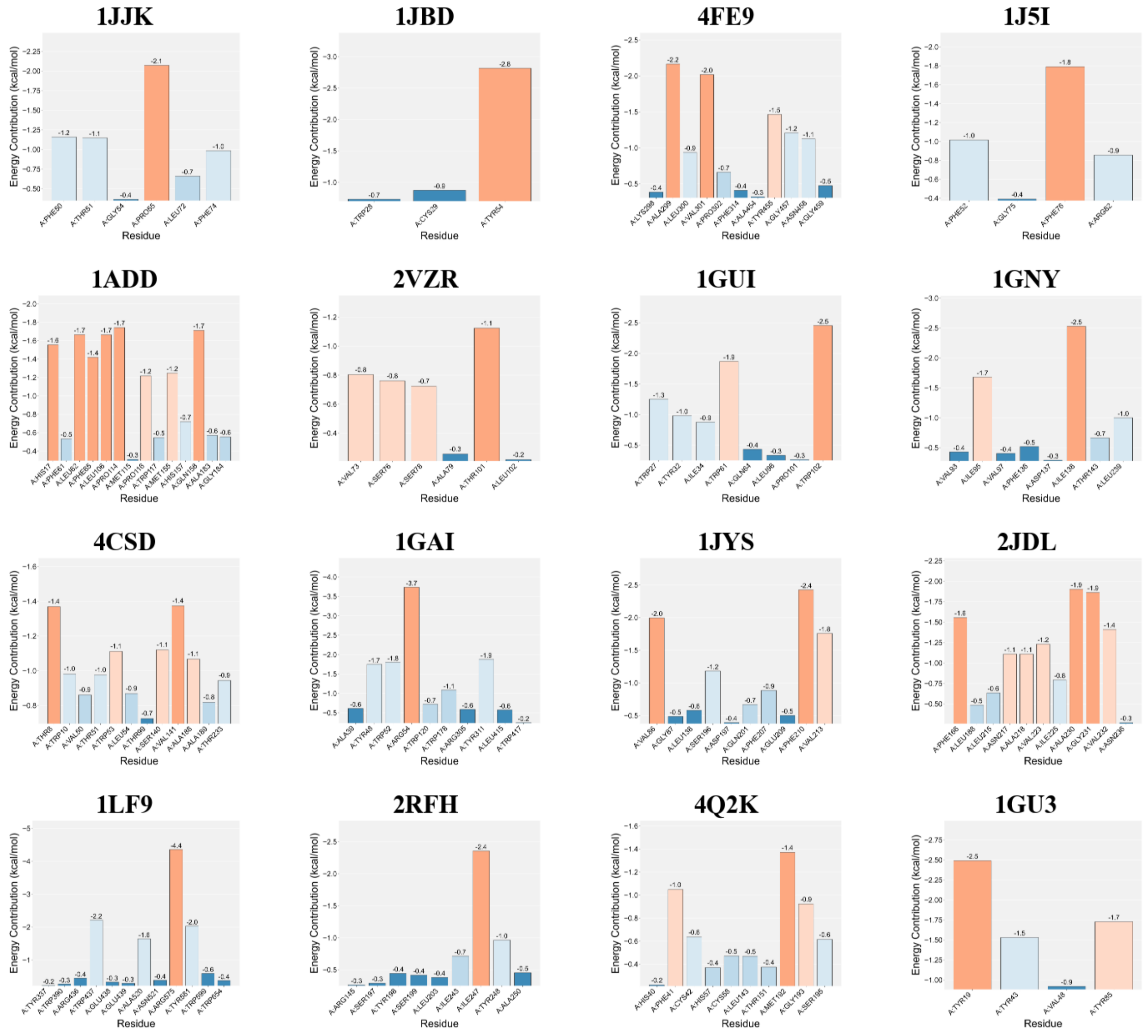


**Fig. S20.** The energy contribution of amino acid residues in the 25(OH)$D_3$ complex with the binding pocket protein.

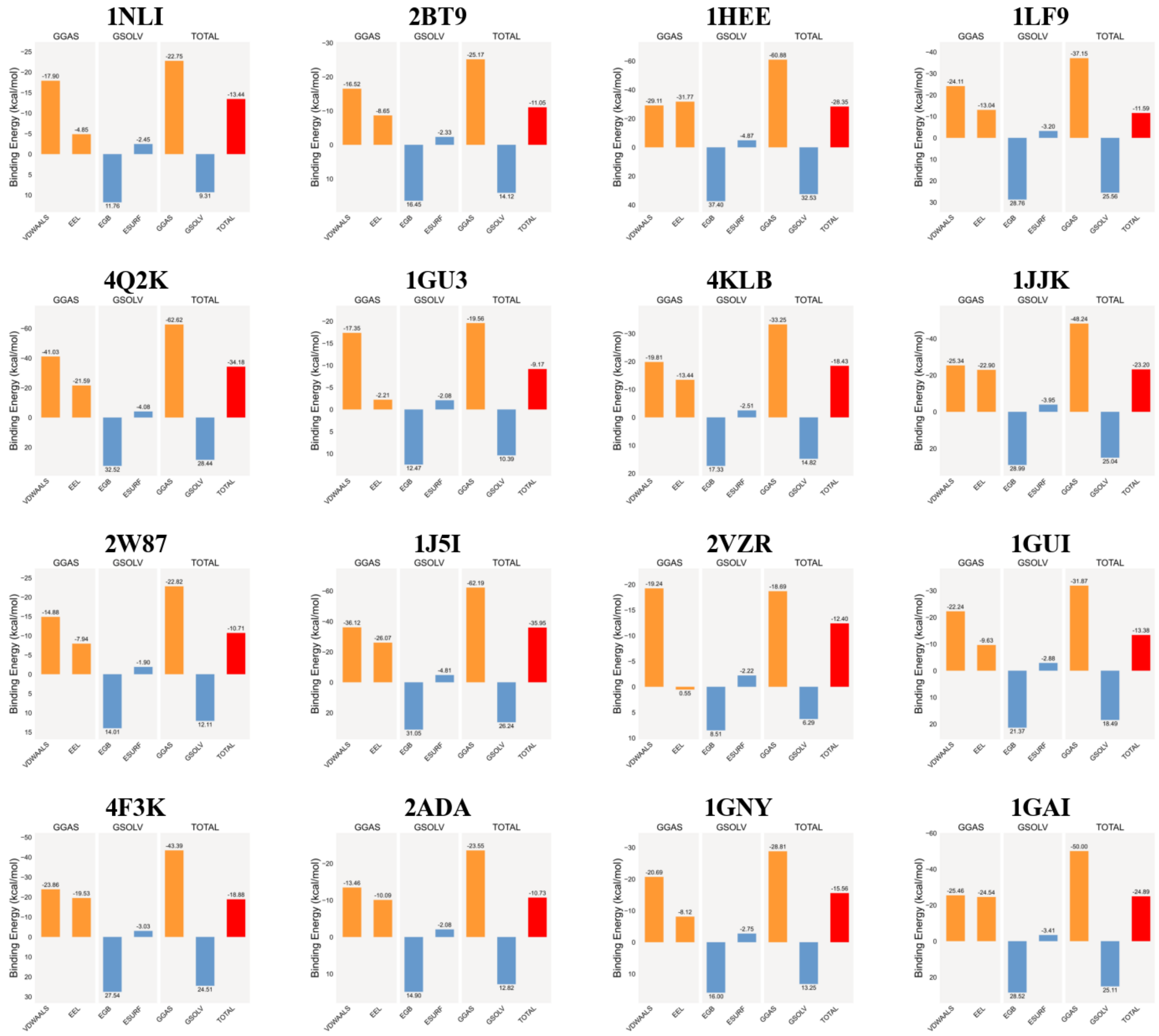


**Fig. S21.** The average binding free energy of Thiamine in complex with the binding pocket protein. VDWAALS, EEL, EGB, ESURF, GGAS, GSOLV, and TOTAL represent van der Waals forces, electrostatic energy, polar solvation energy, nonpolar solvation energy, molecular mechanical energy term, solvation energy term, and the average binding free energy, respectively.

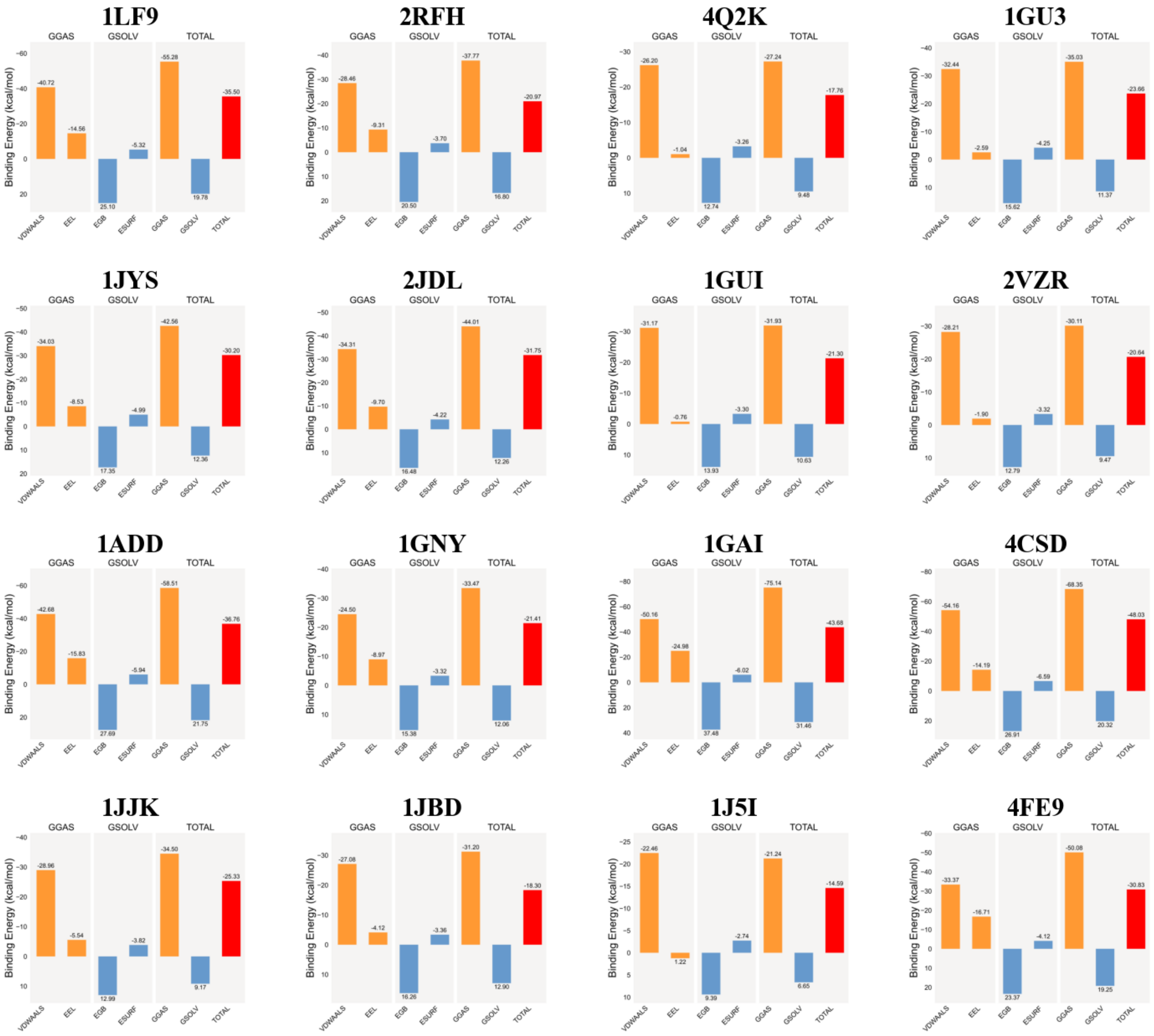


**Fig. S22.** The average binding free energy of 25(OH)$D_3$ in complex with the binding pocket protein. VDWAALS, EEL, EGB, ESURF, GGAS, GSOLV, and TOTAL represent van der Waals forces, electrostatic energy, polar solvation energy, nonpolar solvation energy, molecular mechanical energy term, solvation energy term, and the average binding free energy, respectively.

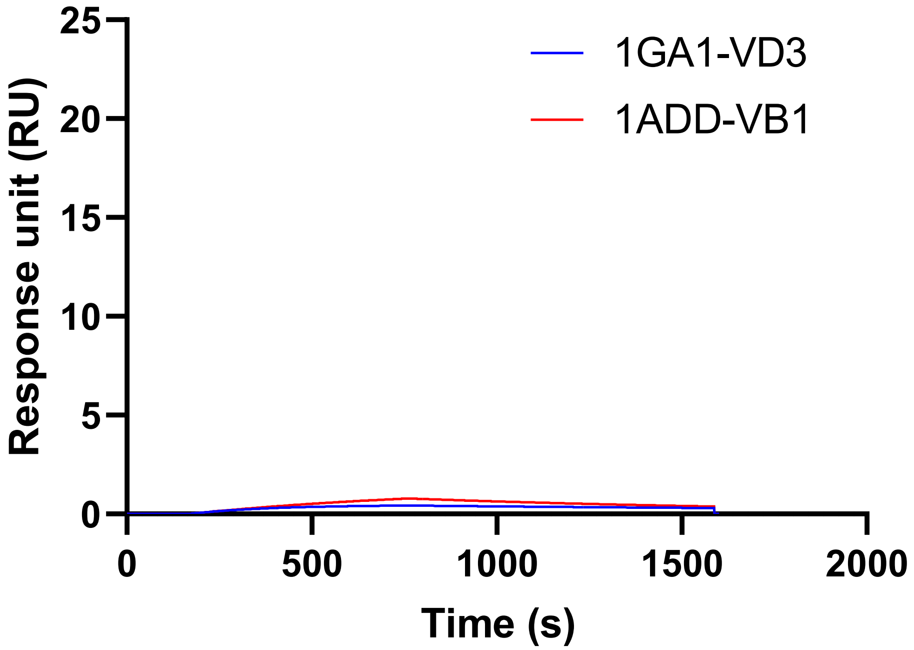


**Fig. S23.** Cross-reactivity testing of 1GAI with $25(OH)D_3$ and 1ADD with Thiamine.

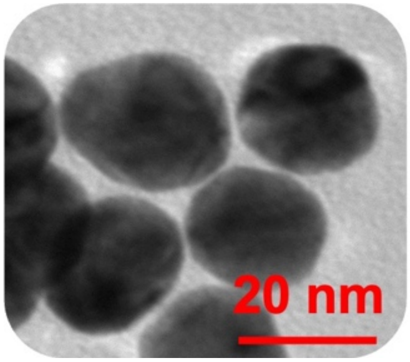


**Fig. S24.** Representative TEM image of the synthesized AuNPs.

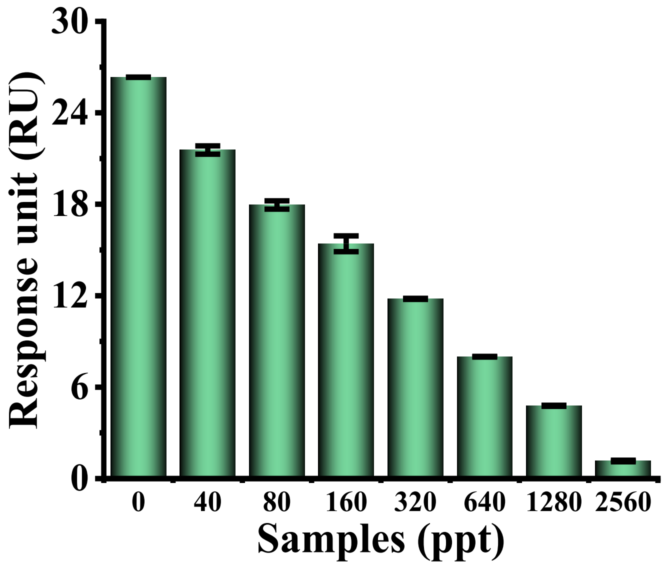


**Fig. S25.** Response results of the MetaSPR sensor chip for Thiamine standard samples

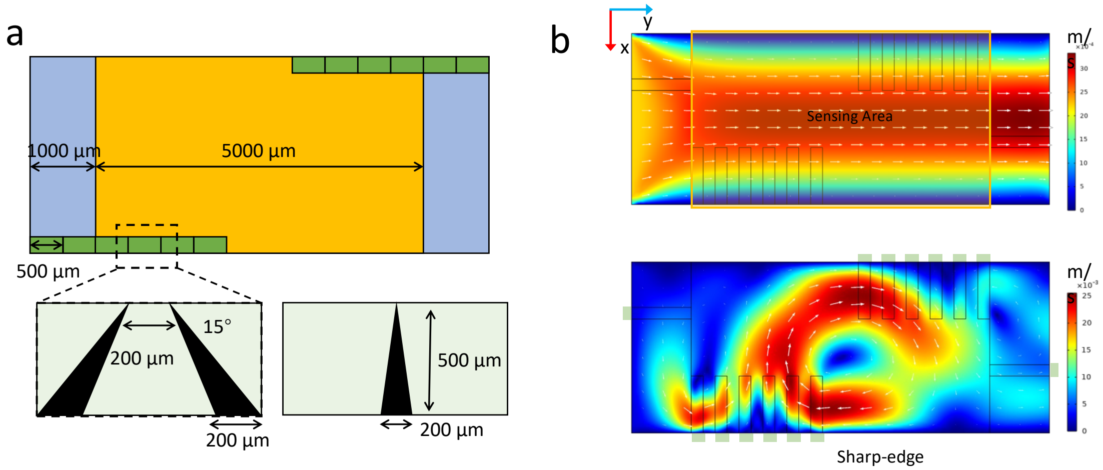


**Fig.S26.** (a) Acoustofluidic sharp-edge array design. (b) Finite element simulation of sharp-edge induced acoustic streaming. 2D velocity magnitude field in the symmetry plane of the microchannel. Streamline visualization showing paired counter-rotating vortices anchored at each sharp edge, covering the entire sensing area.

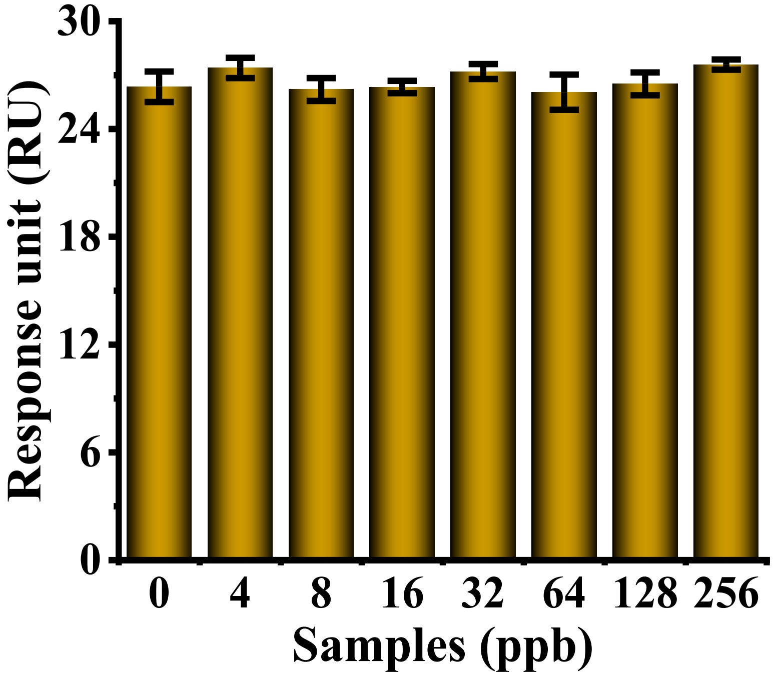


**Fig.S27.** Selectivity test. structurally similar interfering molecules (25(OH)$D_3$, testosterone, thyroid hormone; 256 ppb, 2-fold serial dilution) yielded no significant signal changes

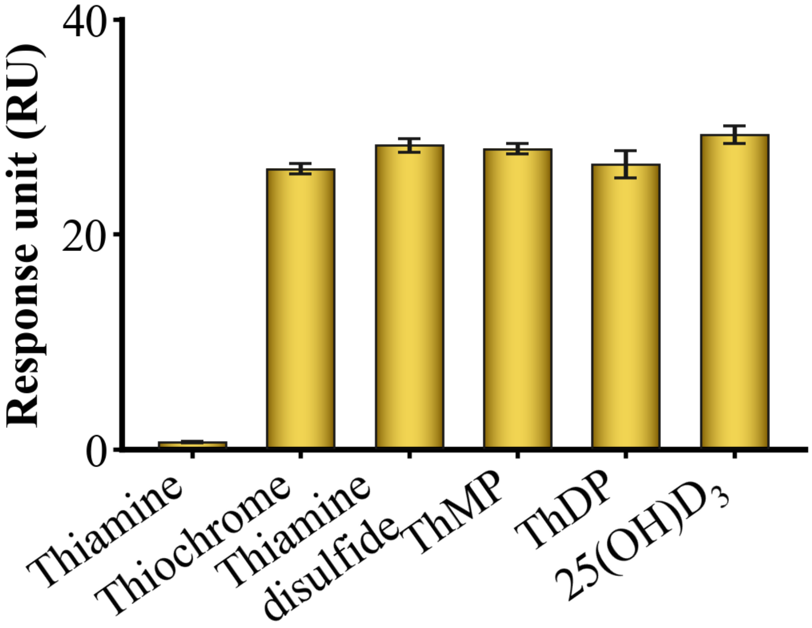


**Fig.S28.** Cross-reactivity assessment of the thiamine assay.

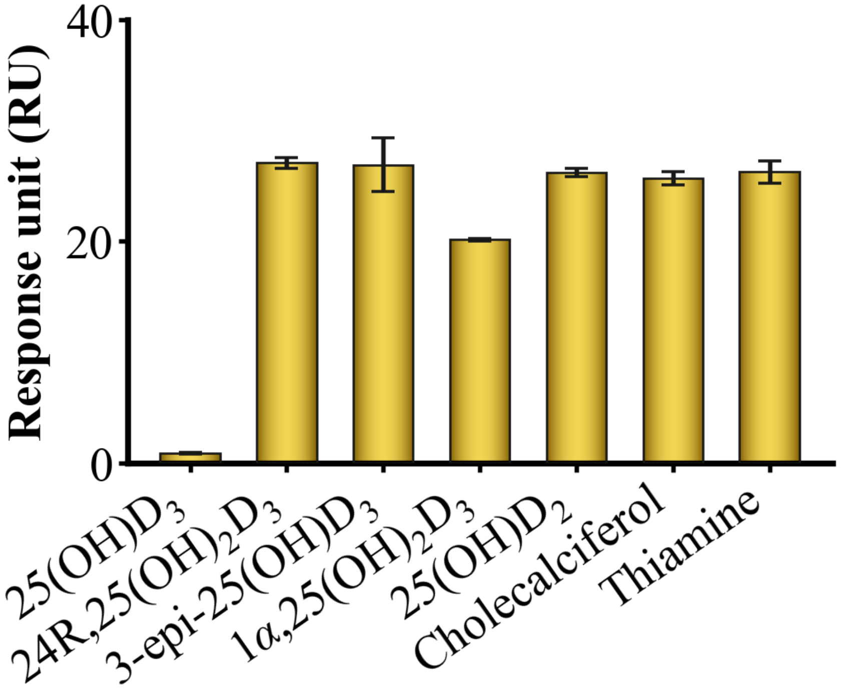


**Fig.S29.** Cross-reactivity assessment of the 25(OH)$D_3$ assay.

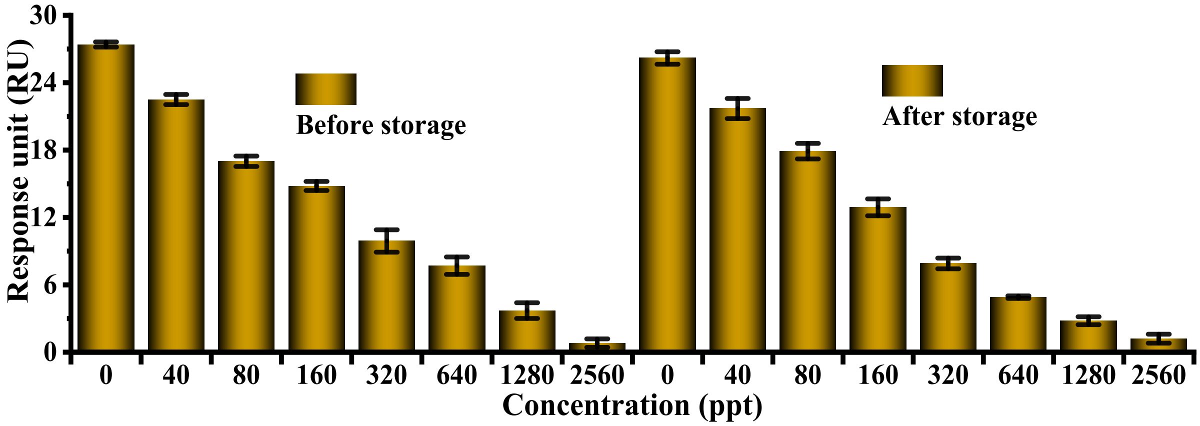


**Fig.S30.** Long-term stability of MetaSPR sensor chip.

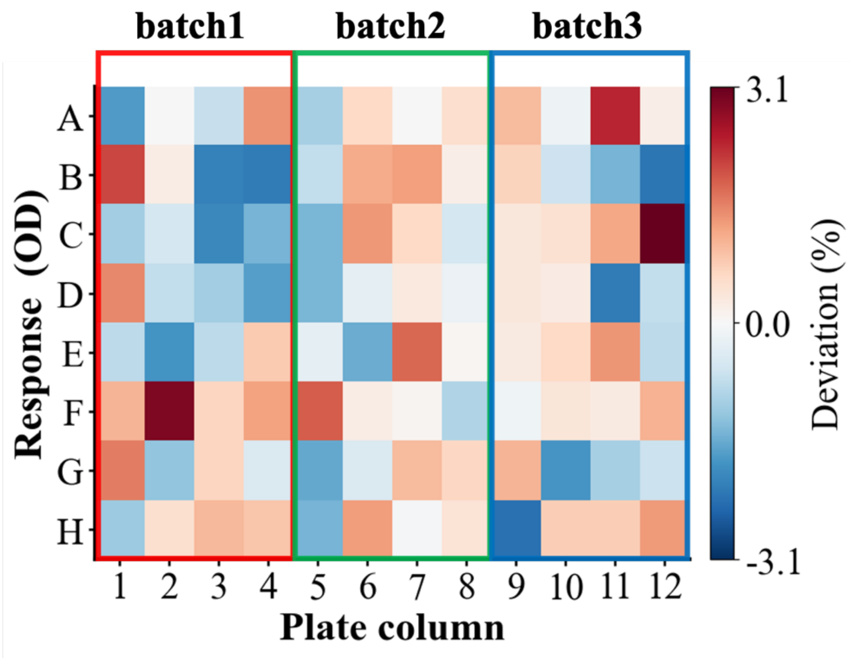


**Fig.S31.** Coefficients of variation of signal responses across chip regions and fabrication batches. Chips from three fabrication batches were integrated into a 96-well plate for evaluation.

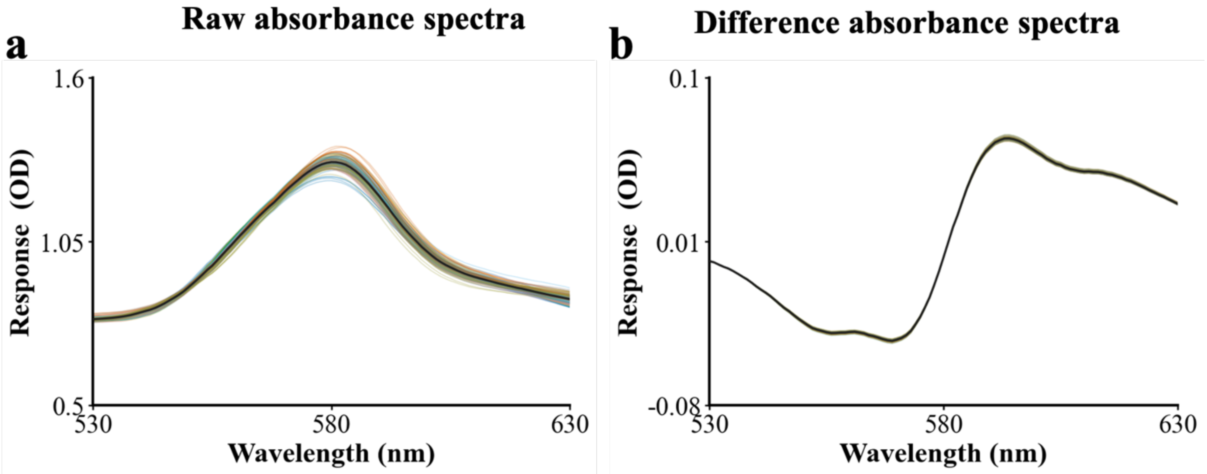


**Fig.S32.** Raw and differential optical density (OD) responses across chip regions and fabrication batches.

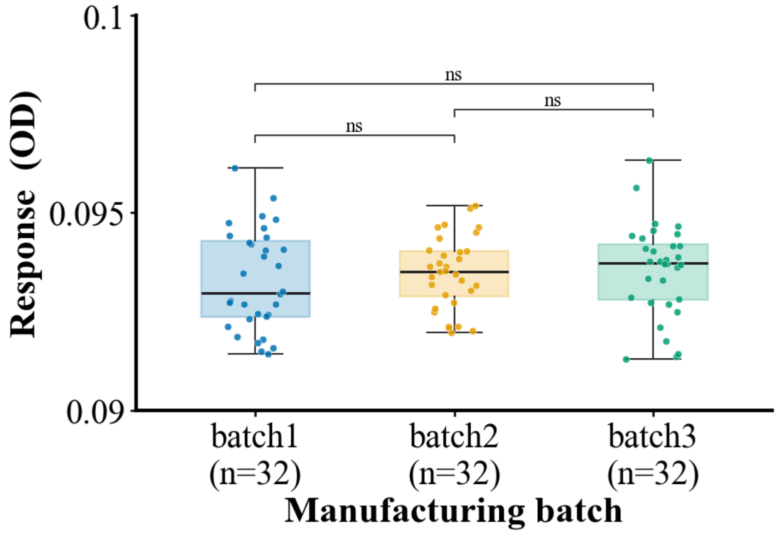


**Fig.S33.** Distribution and uniformity of signal responses across chip regions and fabrication batches.

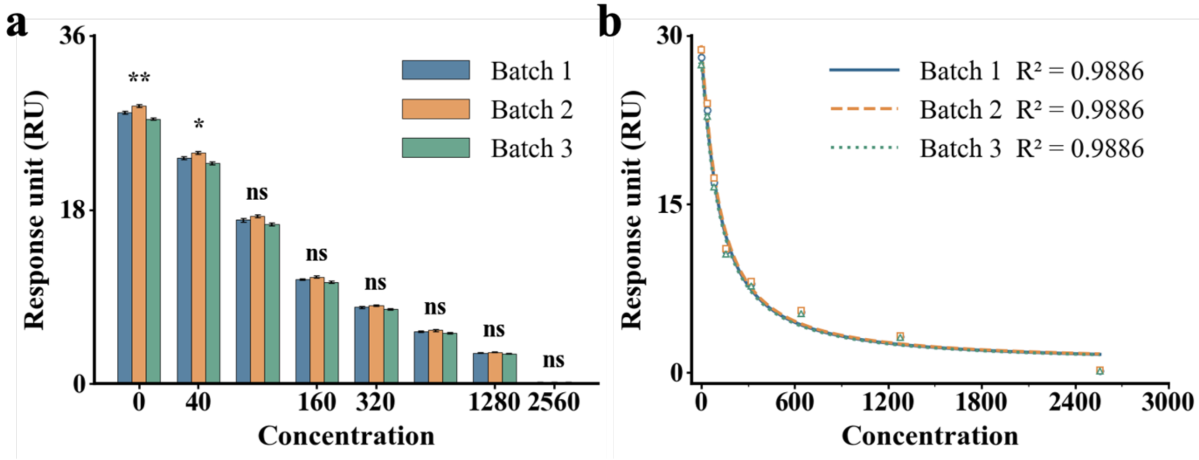


**Fig.S34.** Consistency of thiamine quantification across chip regions and fabrication batches, assessed using calibration curves.

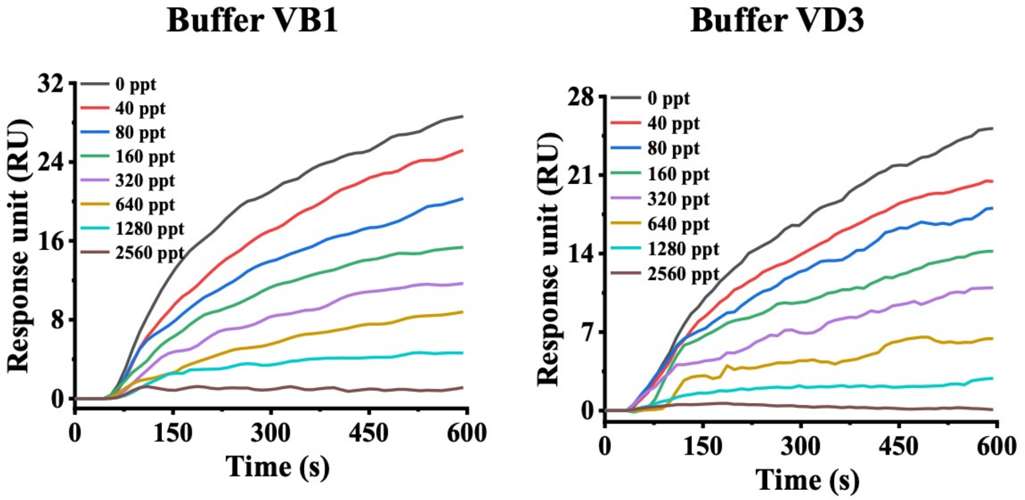


**Fig.S35.** Real-time kinetic curves of the pocket protein biosensor for Thiamine and 25(OH)$D_3$ detection in buffer.

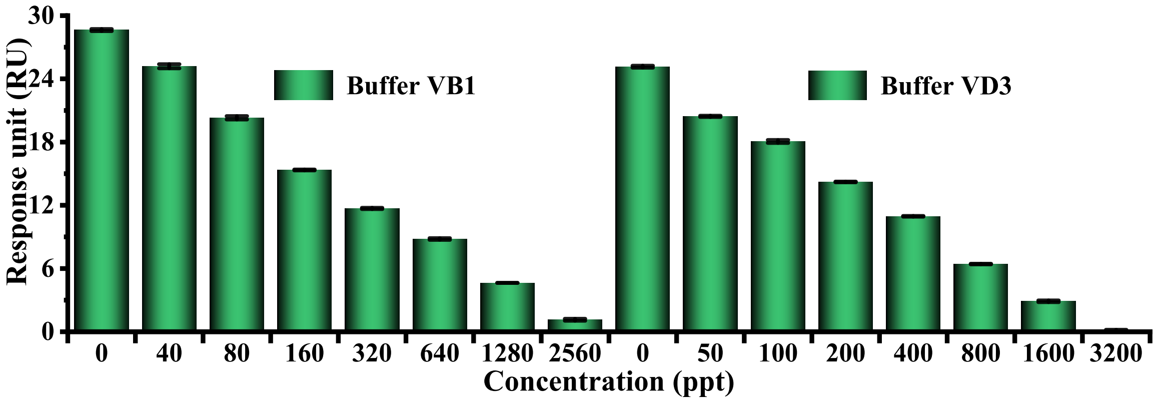


**Fig.S36.** Response units for simultaneous detection of Thiamine and 25(OH)$D_3$ at various concentrations in buffer.

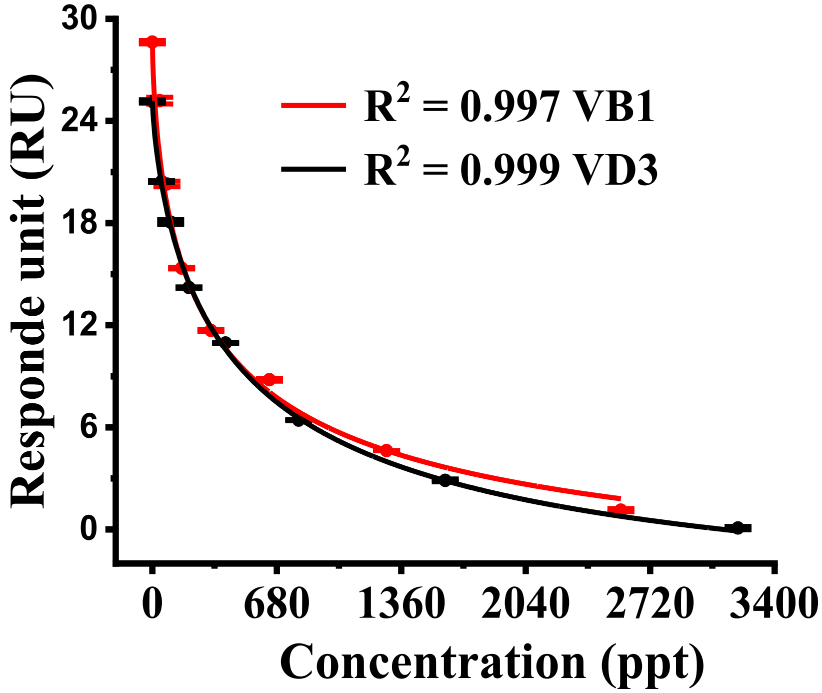


**Fig.S37.** 4-PL fitting curves for one-step detection of Thiamine and 25(OH)$D_3$ in buffer.

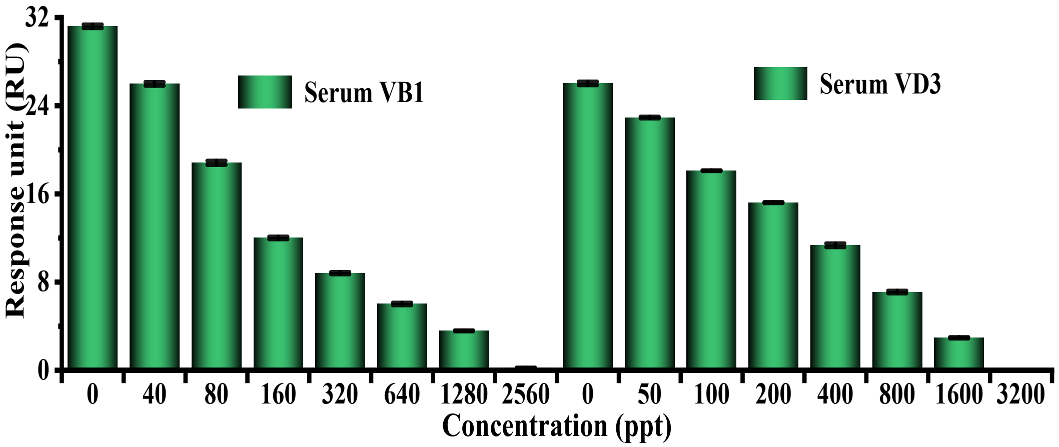


**Fig. S38.** Response signal of the pocket protein biosensor for Thiamine and 25(OH)$D_3$ detection in serum, demonstrating its capability for multiplexed analysis in complex matrices.

**Table S3. Top 40 Thiamine and 25(OH)$D_3$ candidate proteins of screened by PLCA model.**

| Thiamine_Protein ID | Affinity Score | 25(OH)$D_3$_Protein ID | Affinity Score |
|---|---|---|---|
| **1gu3** | 10.59699726 | **1gu3** | 10.31281757 |
| **4f2w** | 10.16633415 | **4f2w** | 10.21533298 |
| **1lf9** | 9.957408905 | **1lf9** | 9.961906433 |
| **4f3k** | 9.841187477 | **4f3k** | 9.923280716 |
| **1gai** | 9.839548111 | **1jys** | 9.914726257 |
| **4csd** | 9.834537506 | **4f3c** | 9.889980316 |
| **1gah** | 9.806289673 | **1nc3** | 9.789022446 |
| **1jys** | 9.801412582 | **1y6q** | 9.743588448 |
| **4f3c** | 9.786499977 | **1y6r** | 9.739207268 |
| **2vzr** | 9.739854813 | **1nc1** | 9.735701561 |
| **1nc3** | 9.732252121 | **1gai** | 9.723989487 |
| **1y6q** | 9.702069283 | **1gah** | 9.695357323 |
| **1agm** | 9.701629639 | **4yml** | 9.647432327 |
| **1y6r** | 9.676404953 | **4wkc** | 9.611533165 |
| **1nc1** | 9.664480209 | **1agm** | 9.591482162 |
| **1nli** | 9.611244202 | **1gny** | 9.55802536 |
| **4yml** | 9.601286888 | **1add** | 9.486537933 |
| **2jdl** | 9.561845779 | **1gui** | 9.468858719 |
| **4wkc** | 9.554825783 | **2ada** | 9.454044342 |
| **1gny** | 9.530344009 | **1fkw** | 9.445796967 |
| **1add** | 9.524628639 | **4q2k** | 9.426605225 |
| **1fkw** | 9.500713348 | **2jdl** | 9.405584335 |
| **2ada** | 9.500490189 | **1a4m** | 9.358101845 |
| **1gui** | 9.454520226 | **2vzr** | 9.353274345 |
| **4q2k** | 9.416372299 | **4csd** | 9.323321342 |
| **1a4m** | 9.405990601 | **2rfh** | 9.309228897 |
| **1j5i** | 9.395941734 | **3kgq** | 9.307146072 |
| **2w87** | 9.277400017 | **2ctc** | 9.303781509 |
| **2bt9** | 9.245034218 | **1nli** | 9.295883179 |
| **2rfh** | 9.221769333 | **4fe9** | 9.277901649 |
| **1jjk** | 9.212238312 | **1j5i** | 9.256350517 |
| **3kgq** | 9.199475288 | **1iy7** | 9.225013733 |
| **2ctc** | 9.173323631 | **1cps** | 9.219407082 |
| **1iy7** | 9.147293091 | **1hee** | 9.217780113 |
| **1hee** | 9.136268616 | **1hdq** | 9.196198463 |
| **7cpa** | 9.135769844 | **1cbx** | 9.193488121 |

| | | | |
|---|---|---|---|
| **6cpa** | 9.135642052 | **1jjk** | 9.180145264 |
| **4klb** | 9.134720802 | **6cpa** | 9.159680367 |
| **1cps** | 9.127919197 | **1jbd** | 9.150238037 |
| **8cpa** | 9.125747681 | **8cpa** | 9.137320518 |

**Table S4. Molecular docking score of Thiamine and 25(OH)$D_3$ candidate proteins.**

| Thiamine_Protein | dock Energy | 25(OH)$D_3$_Protein | dock Energy |
|---|---|---|---|
| **1GAI** | -6.244 | **1ADD** | -8.984 |
| **1GNY** | -5.184 | **1GAI** | -8.357 |
| **1GU3** | -5.518 | **1GNY** | -6.885 |
| **1GUI** | -5.949 | **1GU3** | -7.917 |
| **1HEE** | -6.968 | **1GUI** | -8.276 |
| **1J5I** | -5.834 | **1J5I** | -6.769 |
| **1JJK** | -6.302 | **1JBD** | -6.813 |
| **1LF9** | -7.033 | **1JJK** | -7.456 |
| **1NLI** | -5.474 | **1JYS** | -7.739 |
| **2ADA** | -6.37 | **1LF9** | -8.873 |
| **2BT9** | -5.538 | **2JDL** | -7.412 |
| **2VZR** | -5.611 | **2RFH** | -7.408 |
| **2W87** | -4.832 | **2VZR** | -6.027 |
| **4F3K** | -5.943 | **4CSD** | -8.947 |
| **4KLB** | -5.593 | **4FE9** | -7.086 |
| **4Q2K** | -6.304 | **4Q2K** | -7.368 |

**Table S5. MMPBSA results of Thiamine and 25(OH)$D_3$ candidate proteins.**

| Thiamine_Protein | MD Energy (kcal/mol) | 25(OH)$D_3$_Protein | MD Energy (kcal/mol) |
|---|---|---|---|
| **1GU3** | -9.17 | **1GU3** | -23.66 |
| **1LF9** | -11.59 | **1LF9** | -35.5 |
| **4F3K** | -18.88 | **1JYS** | -30.2 |
| **1GAI** | -24.89 | **1GAI** | -43.68 |
| **2VZR** | -12.4 | **1GNY** | -21.41 |
| **1NLI** | -13.44 | **1ADD** | -36.76 |
| **1GNY** | -15.56 | **1GUI** | -21.3 |
| **2ADA** | -10.73 | **4Q2K** | -17.76 |
| **1GUI** | -13.38 | **2JDL** | -31.75 |
| **4Q2K** | -34.18 | **2VZR** | -20.64 |
| **1J5I** | -35.95 | **4CSD** | -48.09 |
| **2W87** | -10.71 | **2RFH** | -20.97 |
| **2BT9** | -11.05 | **4FE9** | -30.83 |
| **1JJK** | -23.2 | **1J5I** | -14.59 |
| **1HEE** | -28.35 | **1JJK** | -25.33 |
| **4KLB** | -18.43 | **1JBD** | -18.3 |

**Table S6.** Characteristics of the clinical samples.

| Characteristics | Thiamine deficiency status | Healthy | Vitamin D deficiency status | Healthy |
|---|---|---|---|---|
| Number | 1–30 | 31–60 | 1–30 | 31–60 |
| Gender: male, n (%) | 10 (33.3) | 10 (33.3) | 10 (33.3) | 10 (33.3) |
| Age (years) | 18–59 | 17–59 | 20–61 | 18–57 |
